# Pyruvate:ferredoxin oxidoreductase and low abundant ferredoxins support aerobic photomixotrophic growth in cyanobacteria

**DOI:** 10.1101/2021.08.27.457892

**Authors:** Yingying Wang, Xi Chen, Katharina Spengler, Karoline Terberger, Marko Boehm, Jens Appel, Thomas Barske, Stefan Timm, Natalia Battchikova, Martin Hagemann, Kirstin Gutekunst

## Abstract

The decarboxylation of pyruvate is a central reaction in the carbon metabolism of all organisms. Both the pyruvate:ferredoxin oxidoreductase (PFOR) and the pyruvate dehydrogenase (PDH) complex catalyze this reaction. Whereas PFOR reduces ferredoxin, the PDH complex utilizes NAD^+^. Anaerobes rely on PFOR, which was replaced during evolution by the PDH complex found in aerobes. Cyanobacteria possess both. Our data challenge the view that PFOR is exclusively utilized for fermentation. Instead, we show, that the cyanobacterial PFOR is stable in the presence of oxygen *in vitro* and is required for optimal photomixotrophic growth under aerobic conditions while the PDH complex is inactivated under the same conditions. We found that cells rely on a general shift from utilizing NAD(H)-dependent to ferredoxin-dependent enzymes under these conditions.

The utilization of ferredoxins instead of NAD(H) saves a greater share of the Gibbs free energy, instead of wasting it as heat. This obviously simultaneously decelerates metabolic reactions as they operate closer to their thermodynamic equilibrium. It is common thought that during evolution, ferredoxins were replaced by NAD(P)H due to their higher stability in an oxidizing atmosphere. However, utilization of NAD(P)H could also have been favored due to a higher competitiveness because of an accelerated metabolism.

## Introduction

### FeS clusters, pyruvate:ferredoxin oxidoreductase and ferredoxins

Live evolved under anaerobic conditions in an environment that was reducing and replete with iron and sulfur. Later on, hydrogen escape to space irreversibly oxidized Earth (1, 2). Prebiotic redox reactions, that took place on the surfaces of FeS minerals, are at present mimicked by catalytic FeS clusters in a plethora of enzymes and redox carriers (3, 4). One example are ferredoxins, that are small, soluble proteins containing 4Fe4S, 3Fe4S or 2Fe2S clusters and shuttle electrons between redox reactions. They display a wide range of redox potentials between -240 mV to -680 mV and are involved in a variety of metabolic pathways (5). Ferredoxins are among the earliest proteins on Earth and are accordingly present in all three kingdoms of life (6). FeS enzymes are especially widespread in anaerobes (7).

The advent of oxygenic photosynthesis necessitated adaptations, as especially 4Fe4S clusters are oxidized and degraded to 3Fe4S in the presence of oxygen resulting in inactivated enzymes (7–9). In aerobes, FeS enzymes are commonly replaced by FeS cluster free isoenzymes or alternative metabolic strategies (8). One well known example is the replacement of the FeS cluster containing pyruvate:ferredoxin oxidoreductase (PFOR), which catalyzes the decarboxylation of pyruvate during fermentation in anaerobes, by the pyruvate dehydrogenase (PDH) complex for respiration in aerobes (7, 10). Both enzymes catalyze the same reaction, whereat PFOR uses ferredoxin as redox partner and the PDH complex reduces NAD^+^. PFORs are evolutionary old enzymes. They are widespread in autotrophic and heterotrophic bacteria, in archaea, amitochondriate eukaryotic protists, hydrogenosomes as well as in cyanobacteria and algae (7). Depending on organism, metabolism and conditions, PFOR can be involved in the oxidation of pyruvate for heterotrophy or alternatively catalyze the reverse reaction by fixing CO_2_ and forming pyruvate from acetyl CoA for an autotrophic lifestyle (11–13). The enzyme might have played a central role for the evolution of both autotrophic and heterotrophic processes from the very beginning (14). PFOR indeed participates as CO_2_ fixing enzyme in four out of seven currently known and most ancient autotrophic pathways (reverse tricarboxylic acid (rTCA) cycle, reversed oxidative tricarboxylic acid (roTCA) cycle, reductive acetyl-CoA pathway, and dicarboxylate/hydroxybutyrate (DC/HB) cycle) (12, 15). PFORs contain one to three 4Fe4S clusters and get in general readily inactivated by oxygen upon purification. So far, there are only three reported exceptions to this rule: the PFORs of *Halobacterium halobium*, *Desulvovibrio africanus* and *Sulfolobus acidocaldarius* are stable *in vitro* in the presence of oxygen (11, 16–19). Even though all three enzymes are stable upon purification in the presence of oxygen, anaerobic conditions are required for *in vitro* maintenance of enzyme activities with the PFORs of *Desulvovibrio africanus* and *Sulfolobus acidocaldarius.* The enzyme of *Halobacterium halobium* is the only reported PFOR so far, which is active under aerobic conditions *in vitro* (19, 20). *In vivo* studies on these PFORs under aerobic conditions are missing. Ferredoxins that contain 4Fe4S clusters are likewise vulnerable to oxidative degradation. In the evolution from anoxygenic to oxygenic photosynthesis, the soluble 4Fe4S ferredoxin, which transfers electrons from FeS-type photosystems PSI to other enzymes in anoxygenic photosynthesis was replaced by an oxygen-tolerant 2Fe2S ferredoxin (9). In addition, ferredoxins have in general been complemented or replaced by NAD(P)H as alternative, oxygen-insensitive reducing agents in aerobes (10).

### The pyruvate dehydrogenase complex

The PDH complex, which utilizes NAD^+^ is composed of the three subunits: pyruvate dehydrogenase (E1), dihydrolipoyl transacetylase (E2) and dihydrolipoyl dehydrogenase (E3). It catalyzes the irreversible decarboxylation of pyruvate. The PDH complex is active under oxic conditions but gets inactivated under anaerobic conditions in both prokaryotes and eukaryotes, albeit via distinct mechanisms. In the absence of oxygen NADH/NAD^+^ ratios rise as respiration does no longer oxidize the NADH coming from the PDH complex and the subsequent reactions of the TCA cycle. In prokaryotes, as e.g. *E. coli*, NADH interacts with the dihydrolipoyl dehydrogenase (E3) subunit and thereby inhibits the PDH complex (21, 22). In eukaryotes, the PDH complex gets inactivated at high NADH/NAD^+^ ratios via phosphorylation of highly conserved serine residues in the pyruvate dehydrogenase (E1) subunit (23).

*Synechocystis* sp. PCC 6803 is a cyanobacterium that performs oxygenic photosynthesis and lives photoautotrophically by fixing CO_2_ via the Calvin-Benson-Bassham (CBB) cycle. In the presence of external carbohydrates these are metabolized additionally, resulting in a photomixotrophic lifestyle. In darkness *Synechocystis* switches to a heterotrophic or under anaerobic conditions to a fermentative lifestyle. As in many cyanobacteria, pyruvate can be either decarboxylated via PFOR or alternatively via the PDH complex in *Synechocystis.* PFOR is assumed to be involved in fermentation under anoxic conditions and the PDH complex in aerobic respiration. The observation that *pfor* is transcribed under photoautotrophic conditions in the presence of oxygen in the cyanobacteria *Synechococcus* sp. PCC 7942 and *Synechocystis* was therefore surprising but is well in line with the observation that other enzymes assigned to anaerobic metabolism in eukaryotes are expressed in the presence of oxygen as well (10, 24). *Synechoystis* possesses a network of up to 11 ferredoxins containing 2Fe2S, 3Fe4S and 4Fe4S clusters (25, 26). The 2Fe2S ferredoxin 1 (Ssl0020) is essential and by far the most abundant ferredoxin in *Synechocystis* and is the principal acceptor of photosynthetic electrons at PSI (27). Structures, redox potentials and distinct functions have been resolved for some of the alternative low abundant ferredoxins, however, the metabolic significance of the complete network is still far from being understood (25, 26, 28–31).

In this study we show that PFOR and low abundant ferredoxins are required for optimal photomixotrophic growth under oxic conditions. In line with this we found that the cyanobacterial PFOR is stable in the presence of oxygen *in vitro*. PFOR and ferredoxins can functionally replace the NAD^+^ dependent PDH complex, which we found to get inactivated at high NADH/NAD^+^ ratios. Likewise, the ferredoxin dependent F-GOGAT (<u>g</u>lutamine <u>o</u>xo<u>g</u>lutarate <u>a</u>minotransferase) is essential for photomixotrophic growth as well and cannot be functionally replaced by the NADH dependent N-GOGAT. The cells obviously switch in their utilization of isoenzymes and redox pools. However, the key factor for this switch is not oxygen but are the highly reducing conditions within the cells. Our data suggest that the pool of ferredoxins in *Synechocystis* functions as an overflow basin to shuttle electrons, when the NADH/NAD^+^ pool is highly reduced.

## Results

The roles of PDH complex and PFOR were studied in *Synechocystis* under different growth conditions. PDH could not be deleted from the genome indicating that this enzyme complex is essential, whereas *pfor* was knocked out in a previous study (28). In line with this we found that all fully sequenced cyanobacteria contain a PDH complex, which points out its significancs and that 56 % thereof possess a PFOR in addition (Fig. 1S). We unexpectedly found that the *Synechocystis* Δ*pfor* deletion mutant was impaired in its photomixotrophic growth under oxic conditions in continuous light. Growth impairment was typically visible starting around day three to six of the growth experiment (Fig. 1A and 3A). Under photoautotrophic conditions Δ*pfor* grew just as the WT (Fig. 1A). The oxygen concentration in the photomixotrophic cultures was close to saturation around 250 µMol O_2_ throughout the growth experiment (Fig. S2). Studies on the transcription of *pfor* and the alpha subunit of the pyruvate dehydrogenase (E1) of *pdhA* revealed that both genes are transcribed under photomixo- and photoautotrophic conditions (Fig. S3). These observations raised two questions: Why is the PDH complex, which catalyzes the same reaction as PFOR, not able to compensate for the loss of PFOR? And how can PFOR, which is assumed to be oxygen-sensitive, be of physiological relevance in the presence of oxygen?

**Figure 1:**
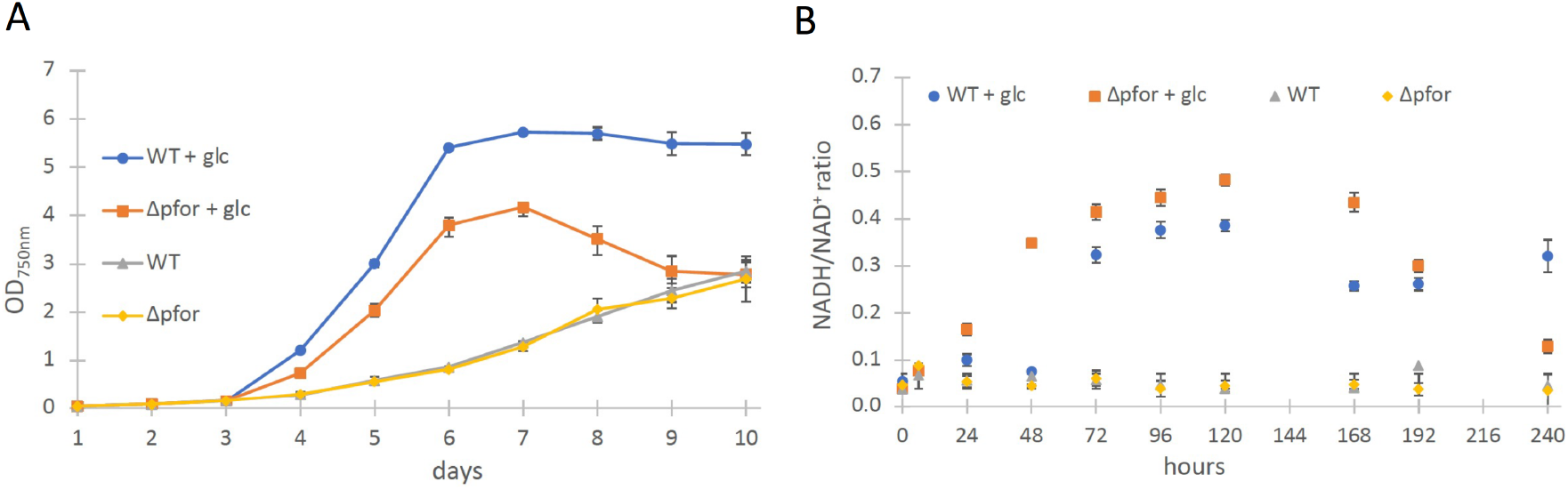
(A) Growth and (B) NADH/NAD^+^ ratios of wild type (WT) and Δ*pfor* under photoautotrophic and photomixotrophic (+ glc) conditions in continuous light. Shown are mean values ± SD from at least 3 replicates.

**Figure 2:**
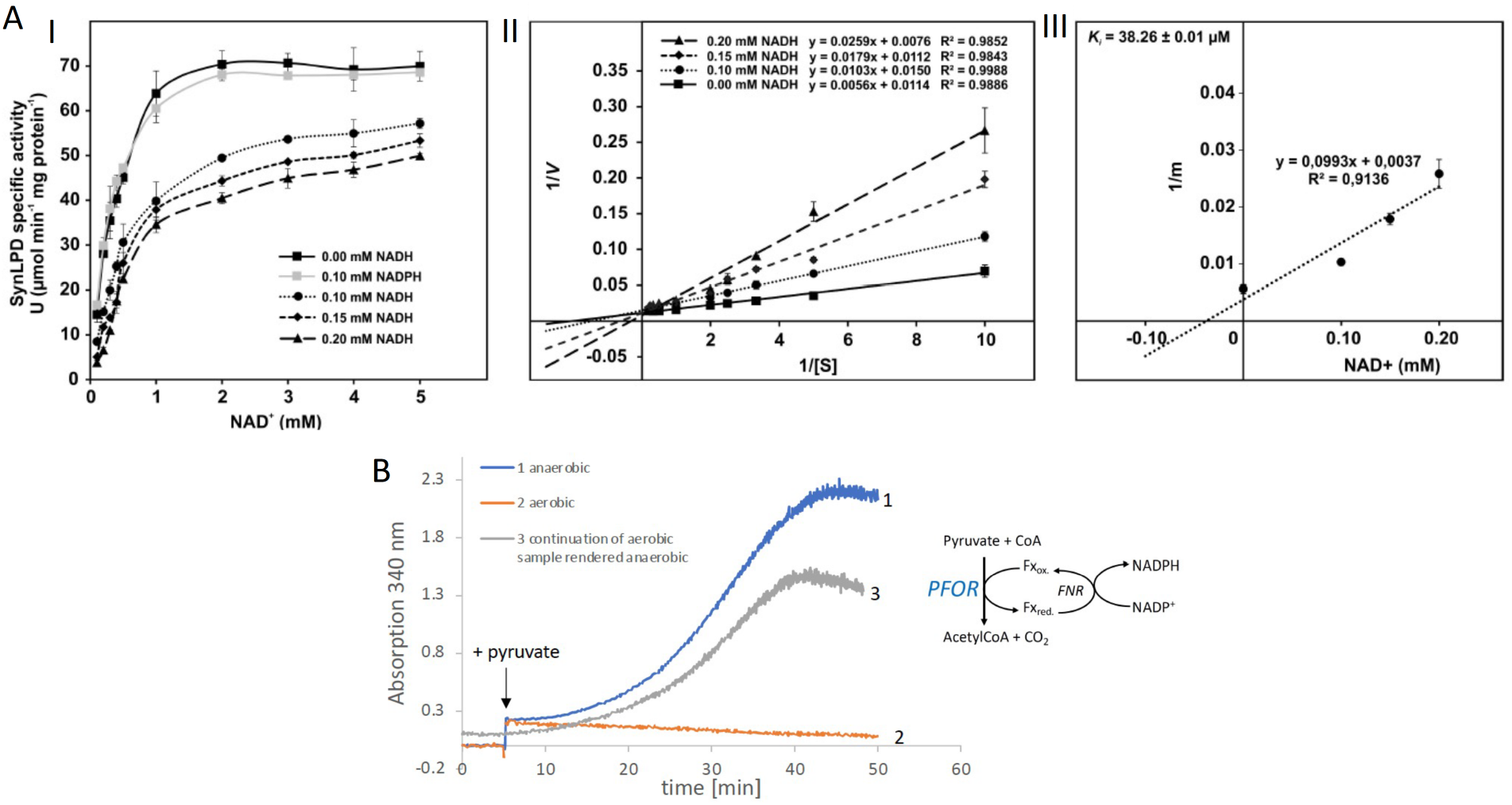
(A) Inhibition of the PDH complex in *Synechocystis* via inactivation of the dihydrolipoyl dehydrogenase (E3) subunit (SynLPD) by NADH. I: The rate of recombinant SynLPD activity (3 mM DL-dihydrolipoic acid) as a function of NAD^+^ (0.1, 0.2, 0.3, 0.4, 0.5, 1, 2, 3, 4 and 5 mM) reduction in the presence of the indicated NADH concentrations (0, 0.1, 0.15 and 0.2 mM). NADPH (0.1 mM) was used as a control to demonstrate the specificity of NADH inhibition. Specific enzyme activity is expressed in µmol NADH per min^−1^ mg protein^−1^ at 25°C. II: Lineweaver-Burk plots of enzyme activities at four NADH concentrations. III: The inhibitor constant (Ki) was estimated by linear regression of (I) the slopes of the three Lineweaver-Burk plots at the four NADH concentrations versus (II) the NADH concentration. Shown are mean values ± SD from at least 3 technical replicates. (B) Enzyme activity of PFOR that was purified in the presence of oxygen. PFOR activity was measured in the presence of FNR, ferredoxin and NADP^+^. The reaction was started by addition of 10 mM pyruvate as indicated by the arrow. Assay 1 (blue line): The assay mixture was kept anaerobic with 40 mM glucose, 40 U glucose oxidase and 50 U catalase, showing that PFOR, which was purified in the presence of oxygen, is active. Assay 2 (red line): Assay 2 had the same composition as assay 1 but glucose, glucose oxidase and catalase were omitted, showing that anaerobic conditions are required for activity of PFOR *in vitro*. Assay 3 (grey line): This assay is the continuation of the measurement of assay 2 after addition of glucose, glucose oxidase and catalase. Representative traces of three replicates are shown.

**Figure 3:**
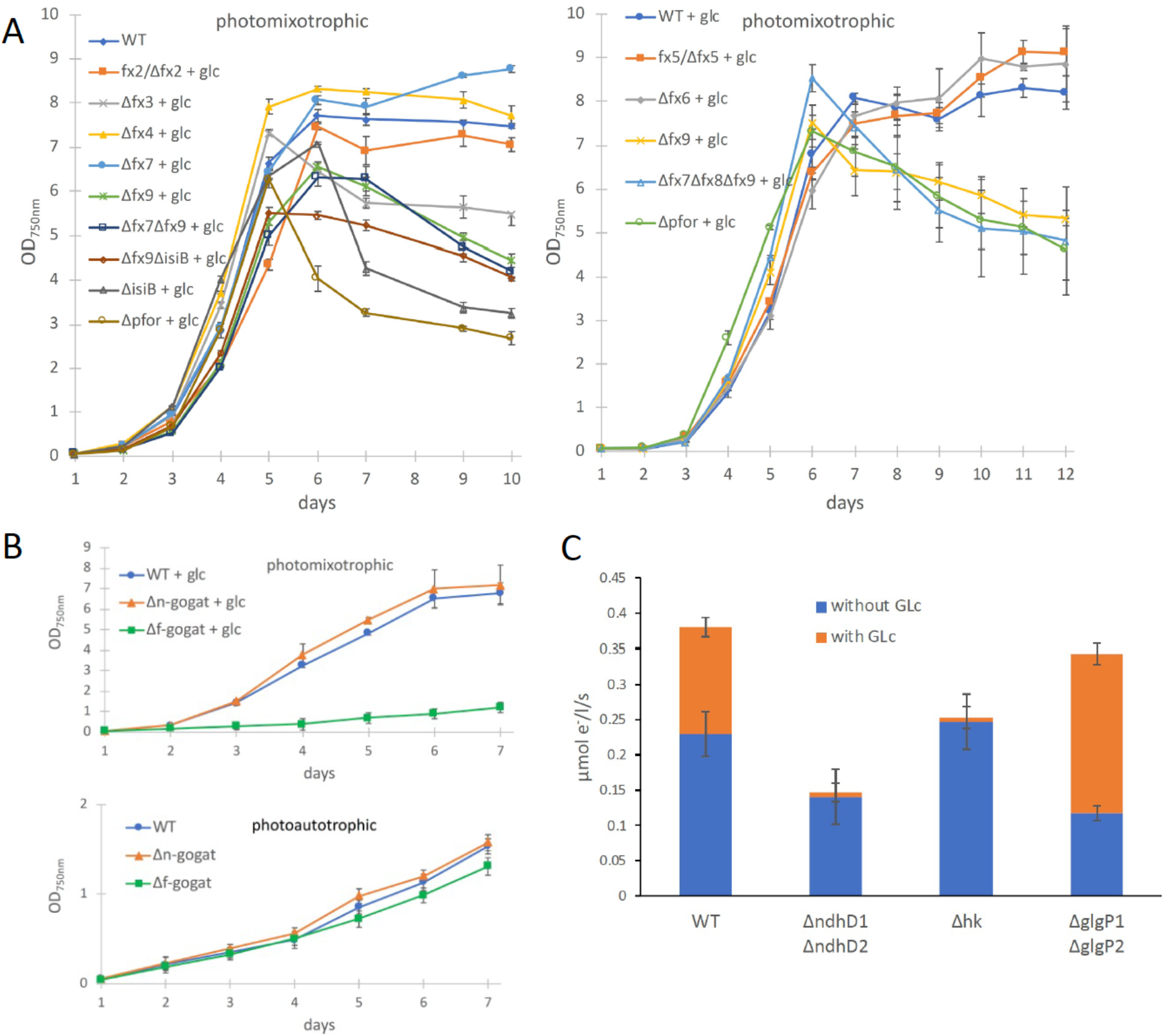
(A) Photomixotrophic growth of wild type (WT), Δ*pfor*, ferredoxin (fx) and flavodoxin (isiB) deletion mutants as indicated. (B) Growth of WT, Δ*f-gogat* and Δ*n-gogat* under photoautotrophic and photomixotrophic conditions. (C) Electron transport with DCMU at PSI in the absence and presence of glucose in the WT, Δ*ndhD1*Δ*ndhD2*, Δ*hk* and ΔglgP1Δ*glgP2*. Shown are mean values ± SD from at least 3 replicates.

The most obvious assumption is that the PDH complex might get inactivated under photomixotrophic conditions. As the PDH complex gets inactivated at high NADH/NAD^+^ ratios in prokaryotes and eukaryotes, we wondered if NADH/NAD^+^ ratios might be increased under photomixotrophic conditions. Corresponding measurements confirmed this assumption. Whereas NADH/NAD^+^ ratios were stable under photoautotrophic conditions in WT and Δ*pfor* they raised three to fourfold in the first five days of photomixotrophic growth, exactly in that period in which the growth impairment of Δ*pfor* in the presence of glucose was most apparent (Fig.1B). In eukaryotes serine kinases phosphorylate three conserved serine residues of the pyruvate dehydrogenase (E1) at high NADH/NAD^+^ ratios and thereby inhibit the PDH complex. In line with this, photomixotrophic growth of two out of ten serine/threonine protein kinase (spk) deletion mutants was affected, which indicates that phosphorylation of enzymes is relevant for optimal photomixotrophic growth in *Synechocystis* (Fig. S4).

In eukaryotes, phosphorylation of serine residues at sites 2 and 3 of the E1 subunit reduces enzyme activity moderately, whereas phosphorylation of the serine residue in site 1 alone completely inhibits the PDH complex (32–34). In order to check if the E1 subunit of *Synechocystis* contains theses conserved serine residues as well, sequence alignments were performed and revealed that serine residues 2 and 3 are absent, whereas the serine residue 1 is present in *Synechocystis* and furthermore highly conserved in the E1 subunit of all 932 cyanobacterial PdhA sequences that were analyzed (Fig. S5 and S6). Immunoblot analyses indicate that the PdhA subunit of the PDH complex might either degrade or might get phospohorylated at high NADH/NAD^+^ ratios (Fig. S7 and S8). However, this could not be shown unambiguously as phosphorylation could not be confirmed by mass spectrometry (Table S3).

For prokaryotes it was shown that the PDH complex is inhibited by a distinct mechanism directly by NADH which binds to the dihydrolipoyl dehydrogenase (E3) subunit of the PDH complex. Therefore, the recombinant dihydrolipoyl dehydrogenase of *Synechocystis* (SynLPD) was tested in an *in vitro* assay with different NADH concentrations. The enzyme indeed loses activity at higher NADH/NAD^+^ ratios, whereas NAPDH has no effect (Fig. 2A). The SynLPD activity was completely inhibited by NADH with an estimated Ki of 38.3 µM (Fig. 2A). Hence, the enzyme activity dropped to approximately 50% at a NADH/NAD^+^ ratio of 0.1 (e.g. at 0.2 mM NADH in the presence of 2 mM NAD^+^). Please note, that much higher NADH/NAD^+^ ratios (> 0.4) were measured in photomixotrophic cells of *Synechocystis* (see Fig. 1B). This poins to an efficient inhibition of PDH activity via the highly decreased function of the E3 subunit (SynLPD). NADH/NAD^+^ ratios above 0.1 could not be tested in the enzyme assays due to the high background absorption of the added NADH, which prevented SynLPD activity detections. Taken together these measurements convincingly show that the PDH complex is most likely inhibited under photomixotrophic conditions at high NADH/NAD^+^ ratios, which provides evidence that pyruvate oxidation must be performed instead via PFOR and gives an explanation for the importance of PFOR under these conditions.

As the cyanobacterial PFOR is regarded as an oxygen sensitive enzyme that exclusively supports fermentation under anaerobic conditions, we overexpressed the enzyme and purified it in the presence of oxygen in order to check for its stability under aerobic conditions (Fig. S9, S10, S11C). Enzymatic tests revealed that PFOR from *Synechocystis* was indeed stable under aerobic conditions *in vitro*, which means that the enzyme was not degraded and kept its activity but required anoxic conditions for the decarboxylation of pyruvate (Fig. 2B) as reported for the oxygen stable PFORs of *Desulvovibrio africanus* and *Sulfolobus acidocaldarius* (*11, 16*).

In contrast to the PDH complex, PFOR transfers electrons from pyruvate to oxidized ferredoxin. In order to investigate if any of the low abundant ferredoxins (Fx) might be of importance for photomixotrophic growth, respective deletion mutants were generated (Table S1 and S2, Fig. S11A) and tested for their ability to grow under photoautotrophic and photomixotrophic conditions. To this end *fx3* (*slr1828*), *fx4* (*slr0150*), *fx6* (*ssl2559*), *fx7* (*sll0662*) and *fx9* (*slr2059*) could be completely deleted from the genome, whereas *fx2* (*sll1382*) and *fx5* (*slr0148*) kept some wild type copies of the genes. We did furthermore not succeed to delete *fx8* (*ssr3184*). Flavodoxin (*isiB; sll0284*), which replaces ferredoxins functionally under Fe-limitation was deleted as well. In addition, the double mutants Δ*fx7*Δ*fx9* and Δ*fx9*Δ*isiB* as well as the triple mutant Δ*fx7*Δ*fx8*Δ*fx9* were generated. Photoautotrophic growth of all these ferredoxin deletion mutants was similar to the WT (Fig. S12). However, under photomixotrophic conditions deletion of either *fx3*, *fx9* or flavodoxin (*isiB*) resulted in a growth behavior that was similar to Δ*pfor* (Fig. 3A).

These results indicate that there might be a general shift to utilize the ferredoxin pool as soon as the NADH/NAD^+^ pool is over reduced. Beside the PFOR/PDH complex couple, GOGAT (<u>g</u>lutamine <u>o</u>xo<u>g</u>lutarate <u>a</u>minotransferase) is as well present in form of two isoenzymes in *Synechocystis* that either utilizes reduced ferredoxin (F-GOGAT; *sll1499*) or NADH (N-GOGAT; *sll1502*). In line with our assumption that ferredoxin utilization is preferred in over reduced cells after glucose addition, we hypothesized that F-GOGAT might be required for optimal photomixotrophic growth. Respective deletion mutants were generated (Table S1 and S2, Fig. S11B) and revealed that neither Δ*f-gogat* nor Δ*n-gogat* were impaired in their growth under photoautotrophic conditions, whereas Δ*f-gogat* displayed a strong growth impairment under photomixotrophic conditions in contrast to Δ*n-gogat* and the WT (Fig. 3B). These data indicate that cells indeed rely on a general switch from utilizing NAD(H) to utilizing ferredoxins for optimal photomixotrophic growth. It was recently shown that photosynthetic complex I (NDH1) exclusively accepts electrons from reduced ferredoxin instead of NAD(P)H (35). Photosynthesis continues under photomixotrophic conditions. However, in addition to water oxidation at photosystem II (PSII), electrons from glucose oxidation can as well enter the respiratory/photosynthetic electron transport chain and eventually arrive at photosystem I (PSI) to support anoxygenic photosynthesis. Anoxygenic photosynthesis thus uses electrons from glucose oxidation that enter the respiratory/photosynthetic electron transport chain and are excited at PSI.

Three entry points exist that can feed electrons from glucose oxidation into the plastoquinone (PQ) pool in the thylakoid membrane: the succinate dehydrogenase (SDH), which accepts electrons from the conversion of succinate to fumarate; NDH-2, which accepts electrons from NADH and photosynthetic complex I (NDH-1), which accepts electrons from reduced ferredoxin (see Fig. 4B). Based on the observed shift from utilizing ferredoxin instead of NAD(P)H, we thus wondered if photosynthetic complex I (NDH-1) might be required for anoxygenic photosynthesis under photomixotrophic conditions as an entry point for electrons coming from glucose oxidation. Cells were incubated with DCMU that blocks the electron transfer from PSII to the PQ-pool. Thereby, exclusively electron transfer from glycogen or glucose oxidation to PSI could be measured based on a recently developed protocol (36). According to this protocol electrons were counted that flow through PSI via DIRK_PSI_ measurements by the KLAS/NIR instrument. The electron transport at PSI was then measured in the absence and in the presence of glucose. In addition to the WT, several mutants were analyzed with deletions in entry points as well as glucose metabolizing enzymes. The mutant with a deleted photosynthetic complex I (Δ*ndhD1*Δ*ndhD2*) should no longer be able to feed electrons from reduced ferredoxin into the respiratory/photosynthetic electron transport chain, while the hexokinase mutant (Δ*hk*) should no longer be able to metabolize external glucose. The glycogen phosphorylase mutant (Δ*glgP1*Δ*glgP2*) is unable to break down its internal glycogen reservoir (36–38). As expected and in parts shown recently (36), addition of glucose resulted in higher donations of electrons to PSI in the WT and Δ*glgP1*Δ*glgP2*, whereas neither Δ*ndhD1*Δ*ndhD2* nor Δ*hk* were able to provide electrons from glucose oxidation to PSI (Fig. 3C). Anoxygenic photosynthesis using glucose oxidation and PSI thus relies on the ferredoxin dependent photosynthetic complex I. In line with this, it was shown earlier that Δ*ndhD1*Δ*ndhD2* is not able to grow in the presence of glucose and DCMU under photoheterotrophic conditions (39).

**Figure 4:**
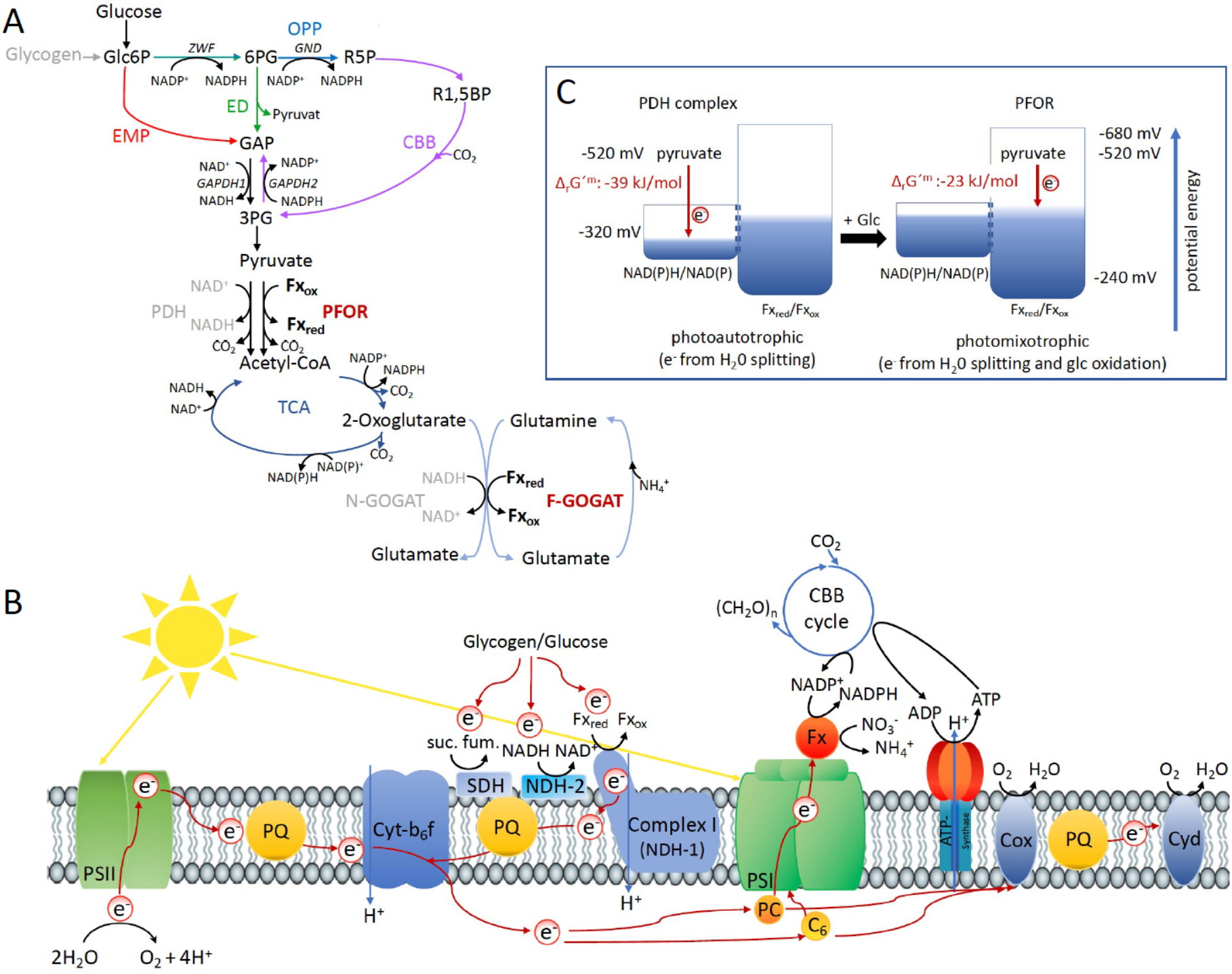
Optimal photomixotrophic growth requires low abundant ferredoxins, PFOR and F-GOGAT. Electrons from glucose oxidation that arrive at PSI require ferredoxin-dependent photosynthetic complex I (NDH-1). Cells shift from utilizing NAD(H) dependent to ferredoxin dependent enzymes when brought from photoautotrophic to photomixotrophic conditions. (A) Glycolytic routes, lower glycolysis and the TCA cycle yield NAD(P)H from glucose oxidation. The only known enzyme that produces reduced ferredoxin from glucose oxidation is PFOR. Both the decarboxylation of pyruvate as well as the synthesis from glutamate from 2-oxoglutarate and glutamine can be catalyzed by distinct enzymes that either utilize ferredoxin (PFOR, F-GOGAT) or NAD(H) (PDH-complex; N-GOGAT). (B) Photosynthetic complex I (NDH-1) accepts electrons from reduced ferredoxin. The complex is required for the input of electrons from glucose oxidation into anoxygenic photosynthesis in the presence of DCMU. (C) The ΔrG’^m^ of pyruvate decarboxylation via the PDH complex is more negative that via PFOR, which results in a higher driving force. Photomixotrophy results in reducing conditions. The redox potential of the NAD(P)H/NAD(P)^+^ pool which is around -320 mV will turn more negative upon reduction. This could facilitate the transfer of electrons from NADH to ferredoxins. In addition, inactivation of NAD^+^ dependent enzymes (such as the PDH complex) and their functional replacement by ferredoxin dependent enzymes (such as PFOR) support the suggested shift from the utilization of the NAD(H) to the ferredoxin pool.

It should be noted in this context that there is no known glycolytic enzyme neither in the Emden-Meyerhoff-Parnass-, the Entner-Doudouroff-, the phosphoketolase-, or the oxidative pentose phosphate pathway in *Synechocystis,* which utilizes ferredoxin as electron acceptor (38, 40, 41). PFOR is currently indeed the only known enzyme in the central carbon metabolism that reduces ferredoxin upon glucose oxidation (Fig. 4A). The second known source for reduced ferredoxin is PSI (Fig.4B). Further studies are required to elucidate the exact role of ferredoxins under photomixotrophic conditions.

## Discussion

There is an irritating high number of reports in prokaryotes and eukaryotes on the presence and expression of enzymes under oxic conditions that are assigned to anaerobic metabolism (10, 24, 42). One example is the production of hydrogen by the oxygen sensitive FeFe-hydrogenase in air-grown *Chlamydomonas reinhardtii* algae in a fully aerobic environment, which is enabled by microoxic niches within the thylakoid stroma (43). Another example is the constitutive expression of PFOR and the oxygen sensitive NiFe-hydrogenase under oxic conditions in cyanobacteria. By itself, the widespread presence of these enzymes in organisms that either live predominantly aerobically as e.g. cyanobacteria or are even obligate aerobes as e.g. *Sulfolobus acidocaldarius*, which possesses a PFOR, indicates a misconception and lack of understanding. The PFOR of *Sulfolobus acidocaldarius* could be isolated as stable enzyme in the presence of O_2_, however, enzyme activity measurements required the consumption of oxygen *in vitro* (11). Does this mean, that anaerobic micro-niches are required within this obligate aerobe to activate an enzyme of its central carbon metabolism? It might alternatively be that living cells have the ability to maintain reducing conditions in the presence of oxygen, which is a challenge in enzymatic *in vitro* assays. Conclusions on *in vivo* enzyme activities based on *in vitro* experiments therefore should be made with caution. Even though we could measure decarboxylation of pyruvate via PFOR only in the absence of oxygen *in vitro*, our data strongly indicate that this enzyme is active *in vivo* under aerobic and highly reducing conditions. We assume that either anaerobic micro-niches or alternatively mechanisms within the cell that are not understood yet, keep the enzyme active in an aerobic environment.

The replacement of FeS enzymes and ferredoxins by FeS free alternatives and NADPH in the course of evolution is in general discussed with regard to the oxygen sensitivity of FeS clusters in connection with the shift from anoxic to oxic conditions on Earth (8, 10). Oxygen is without any doubt one important factor. However, the shift from anoxic to oxic conditions went along with a shift from reducing to more oxidizing conditions. This shift was among others achieved by the escape of hydrogen into space, which irreversibly withdrew electrons from Earth (2). The withdrawal of electrons and the establishment of oxidizing conditions might have been an additional important factor (independent of oxygen and the oxygen sensitivity of FeS clusters) that triggered these evolutionary changes by enabling reactions with higher driving forces. The idea is thus that PFOR and ferredoxins might have been replaced by the PDH complex and NADH due to their potential to release larger amounts of Gibbs free energy (ΔG<<0). When competing with other organisms for resources an accelerated metabolism can be highly beneficial. The decision to either utilize the PDH complex or alternatively PFOR and along this line, the replacement of PFOR by the PDH complex in the course of evolution might have been determined by the prioritization for high chemical driving forces. On that note, we were unable to delete the PDH complex in *Synechocystis,* which points to its essential role. PFOR is in contrast dispensable under photoautotrophic conditions and cells obviously prefer to decarboxylate pyruvate via the PDH complex under these conditions. By transferring electrons to NAD^+^ instead of ferredoxin less Gibbs free energy is stored. However, this comes along with a higher driving force that is visible when regarding the reaction Gibbs energies of Δ_r_G’^m^ -39 kJ/mol for the reaction catalyzed by the PDH complex versus Δ_r_G’^m^ -23 kJ/mol for the reaction catalyzed by PFOR (Fig. 4C) (44).

Under photomixotrophic conditions, photosynthesis and glucose oxidation operate in parallel causing highly reducing conditions in the cells. As soon as the NADH/NAD^+^ pool is highly reduced, cells might be forced to switch to the ferredoxin pool, as shown by our data. Glucose is alternatively oxidized via four glycolytic routes. Flux analyzes have shown that glycolytic intermediates enter the CBB cycle, eventually lower glycolysis and finally provide pyruvate (45). Depending on the precise route taken, glucose oxidation yields distinct forms of reducing equivalents (38). Three enzymes are involved in oxidation steps: Glc6P dehydrogenase (Zwf) and 6PG dehydrogenase (Gnd) yield NADPH, whereas GAP dehydrogenase (GAPDH) yields NADH. NAD(P)H is furthermore provided downstream in the TCA cycle. PFOR is thus the only known direct source for reduced ferredoxin in glucose oxidation (Fig. 4A). The wide network of low abundant ferredoxins in *Synechocystis* and the importance of these ferredoxins under photomixotrophic conditions on the one hand and the low number of known enzymes that directly reduce ferredoxins on the other hand unveils that our conception is not yet inherently consistent. An additional potential source of reduced ferredoxin could be the transfer of electrons from NAD(P)H. The transhydrogenase (PntAB), which is located in the thylakoid membrane utilizes proton translocation to transfer electrons from NADH to NADP^+^ (46). Electrons from NADPH could be further transferred to ferredoxin via ferredoxin-NADPH-oxidoreductase (FNR). Another potential turntable for the exchange of electrons is the diaphorase part of the NiFe-hydrogenase in *Synechocystis*, which was recently shown to shuttle electrons between NAD(P)H, flavodoxin and several ferredoxins *in vitro* (26).

In order to get a complete picture, it would be essential to know the redox potentials of all ferredoxins in *Synechocystis*. Currently, they have been determined for Fx1 (-412 mV), Fx2 (-243 mV), and Fx4 (-440 mV), whereas the value for Fx4 is based on measurements of a homologue in *Thermosynechococcus elongatus* (27, 29, 30). Fx1 to Fx6 in *Synechocystis* possess 2Fe2S clusters for which redox potentials between -240 to -440 mV are typical (5). For 3Fe4S clusters as present in Fx8 (containing one 3Fe4S and one 4Fe4S cluster) redox potentials between -120 to -430 mV were determined and for 4Fe4S clusters as present in Fx7 (4FeFS) and Fx9 (containing two 4Fe4S clusters) redox potentials between -300 to -680 mV were found (5). Without yet knowing the exact values for all ferredoxins in *Synechocystis*, it is anyway likely that they span a wide range of redox potentials.

The redox potential of any given couple does not have a constant value but is influenced among others by the ratio of the redox partners. The redox potential of the NADH/NAD^+^ pool will thus turn more negative upon reduction. Obviously, storing electrons as reducing equivalents with lower redox potential saves more of their potential energy. The driving force of a reaction is on the other hand higher if electrons are transferred across larger redox potential differences. This will slow down the back reaction and thereby speed up the forward reaction. The idea is thus that the NADH/NAD^+^ pool gets reduced first prioritizing high driving forces. However, as the redox potential of the NADH/NAD^+^ pool turns slowly more negative, it might reach levels that are characteristic for ferredoxin couples. This might provoke a metabolic shift to transfer electrons to oxidized ferredoxin instead of NAD^+^ (Fig. 4C). This shift can be regulated on several levels. Among others, as shown in this study, high NADH/NAD^+^ ratios can inactivate enzymes that rely on this couple and thereby support the action of isoenzymes that interact with the Fx_red_/Fx_ox_ couple instead. In addition, electron turntables as the transhydrogenase, FNR and the diaphorase can support this shift (26, 46).

In the case of *Synechocystis*, it is especially beneficial to shift their reducing equivalent pools to ferredoxin, as Fx is able to donate electrons to the photosynthetic complex I (35). In contrast to SDH and NDH-2, the photosynthetic complex I is coupled to a proton gradient and thus yields a higher amount of ATP. By shifting their pools of reducing equivalents, cells are thus able to save a greater share of the potential energy of electrons instead of wasting it as heat. As a pay-off, this shift should obviously come along with a slowdown of metabolic reactions.

## Conclusion

The cyanobacterium *Synechocystis* encounters highly reducing conditions under photomixotrophy in the presence of oxygen. The PDH complex gets inactivated under these conditions at high NADH/NAD^+^ ratios and is functionally most likely replaced by PFOR. PFOR is stable in the presence of oxygen *in vitro* and reduces ferredoxin instead of NAD^+^. PFOR, low abundant ferredoxins and the ferredoxin-dependent GOGAT are required for optimal photomixotrophic growth. Electrons from the oxidation of external glucose furthermore rely upon the presence of photosynthetic complex I (which accepts electrons from ferredoxin) in order to reach PSI. These findings indicate that cells perform a general shift in the utilization of their reducing equivalent pools from NAD(H) to ferredoxin under photomixotrophic conditions.

## Materials and Methods

### Growth conditions

All strains were grown in 50 ml BG-11 (47) buffered with TES pH 8. WT, Δ*pfor*, Δ*f-gogat*, Δ*n-gogat*, Δ*isiB*, all ferredoxin deletion mutants, Δ*ndhD1*Δ*ndhD2*, Δ*hk*, and Δ*glgP1*Δ*glgP2* were and placed in 100 ml Erlenmeyer flasks on a rotary shaker at 28 °C, 50 µE m^−2^ s^−1^ and 100 rpm. After several days of growth, the cells were inoculated into 200 ml BG-11 at an OD_750_ of 0.05 and placed into glass tubes bubbled with air at 50 µE m^−2^ s^−1^ at 28 °C and growth was monitored by measuring the optical density at 750 nm. In liquid cultures all the strains were grown without addition of antibiotics and for photomixotrophic conditions 10 mM glucose was added. In case of the mutants deficient in Serine/Threonine- and Tyrosine kinases all strains were first inoculated from agar plates into 100 ml BG-11-Medium. The cells were pre-cultivated shaking Erlenmeyer flasks (140 rpm) for 7 days under ambient air with 35 µE m^−2^ s^−1^ and 30°C. Ahead of the growth experiment, cells were harvested from the cultures by centrifugation (5000 xg, 20°C, 5 min) and re-suspended in fresh BG11 medium. The cultures were adjusted to OD_750_ of 0.5 with BG-11 medium and subsequently split into two aliquots. One part of the cultures was supplemented with 10 mM Glucose and the remaining part served as a control without glucose. *Synechocystis* strains were grown under ambient air conditions with 35 µE m^−2^ s^−1^ and 30°C. Growth was monitored by measuring the OD_750_ every 24 h until the cultures reached the stationary growth phase. Pictures displaying the phenotypes were taken at the end of the cultivation period.

For mutant selection and seggregation the cells were grown on BG-11-agar containing 50 µg/mL kanamycin, 20 µg/mL spectinomycin, 25 µg/mL erythromycin, 10 µg/mL gentamycin, and 20 µg/mL chloramphenicol.

### Construction of mutants

All the primers used in this study are listed in table S1. All mutants are listed in table S2. Most mutants were constructed in the non-motile GT WT of *Synechocystis* sp. PCC 6803 while a few serine/threonine protein kinase mutants were constructed in the motile PCC-M WT strain of *Synechocystis* as indicated in detail in table S2 (48). The procedure to generate the constructs for deletion of *pfor*, *pdhA*, *isiB* and the different ferredoxin genes was described in Hoffmann et al. (2006) (49). In brief, the up- and downstream regions as well as the required antibiotic resistance cassette were amplified by PCR. Subsequently, the three fragments were combined by a PCR fusion including the outermost primers. The resulting product was inserted by TA-cloning into the pCR2.1 TOPO-vector (ThermoFisher, Waltham, MA, USA). Constructs for the deletion of the genes of the NADH-dependent and the ferredoxin-dependent GOGAT were generated by Gibson cloning (50) assembling three fragments into the pBluescript SK(+) in a single step. After examination by sequencing the plasmids were transformed into *Synechocystis* sp. PCC 6803 cells as described (51). Resulting transformants were either checked by PCR or Southern hybridization after several rounds of segregation (Fig. S11).The deletion strains of the serine/threonine kinase genes *spkA*, *spkB*, *spkD*, *spkG* and *spkL* carry mutations in their kinase domains and were generated accordingly. Their segregation was complete and will be described elsewhere (Barkse and Hagemann, in preparation).To generate a construct for overexpression of *pfor* (*sll0741*) including a His-tag a DNA fragment containing 212 bp up- and 212 bp downstream of the *sll0741* start codon, with a BamHI, XhoI and NdeI site in between and 20 bp sequences that overlap with the pBluescript SK(+) vector at the respective ends was synthesized by GeneScript (Psicataway Township, NJ, USA)(Fig S6A). Another DNA fragment containing a modified petE promotor, followed by His-tag, TEV cleavage recognition site and linker encoding sequences, various restriction sites and 20 bp sequences that overlap with the pBluescript SK(+) vector at the respective ends was also synthesized by GenScript (Fig S6B). These fragments were cloned into the pBluescript SK(+) vector by Gibson cloning, respectively. A kanamycin antibiotic resistance cassette was inserted into the EcoRV site of the plasmid containing the modified petE promotor. The resulting promoter-cassette plasmid and the PFOR plasmid were digested with BamHI and NdeI and the promoter cassette was ligated into the alkaline phosphatase treated PFOR plasmid to yield the final construct. This plasmid was sequenced, transformed into *Synechocystsis* sp. PCC 6803 and segregation was confirmed by PCR analysis (Fig S9C).

### Southern-Blotting

200 ng genomic DNA was digested with Hind III and loaded on a 0.8 % agarose gel in TBE buffer. After blotting the DNA on a nylon membrane (Hybond N+, Merck, Darmstadt, Germany) it was cross-linked to the membrane in a Stratalinker (Stratagene, CA, USA). Detection of the respective bands was carried out by the Dig DNA labeling and detection kit (Roche, Penzberg, Germany) according to the manufacturers instructions.

### RT-PCR

To a volume of 15 μl containing 1 μg of RNA 2 μl RNase-free DNase (10 U/μl, MBI Fermentas, St. Leon-Rot, Germany), 2 μl 10 x DNase buffer (MBI Fermentas, St. Leon-Rot, Germany) and 1 μl Riboblock RNase Inhibitor (40 U/μl, MBI Fermentas, St. Leon-Rot, Germany) were added before incubation at 37 °C for 2 hours. Subsequently the sample was quickly cooled on ice. 2 μl 50 mM EDTA was added and it was incubated at 65 °C for 10 min and again quickly cooled on ice to get rid of the DNase activity. To check the digestion efficiency, 1 μl of the sample was used as a template for PCR. 1 μl genomic DNA and 1 μl H_2_O were used as positive and negative controls, respectively. Reverse transcription PCR was performed with 9 µl of those samples free of DNA with the RT-PCR kit (Applied Biosystems, Karlsruhe, Germany) according to the manufacturer’s instruction. 9 µl of the same sample was used in parallel as a negative control. The reaction mixture was incubated for 1 h at 37 °C including a gene-specific tag-1 primer. For the subsequent PCR a gene-specific tag-2 primer and the respective reverse primer (s. table S1) were used.

### Oxygen measurements

To measure the concentration of dissolved oxygen in the cultures oxygen sensors from Unisense (Unisense, Aarhus, Denmark) were used. After a two point calibration of the sensor by using distilled water equilibrated with air and a solution with 0.1 M NaOH and 0.1 M ascorbate containing no oxygen it was placed in the respective culture and the measurement was started.

### Mass spectrometry

Pieces of the gel corresponding to bands 1-4 in the Fig. 2A were excised, reduced and alkylated following by digestion with Trypsin Gold (Promega) and extraction of peptides as described (52). Peptides from the bands 2-4 were further enriched for phosphopeptides using TiO_2_ as described (53). Next, the peptides were analyzed by LC−MS/MS using a Q Exactive Hybrid Quadrupole-Orbitrap mass spectrometer (Thermo Scientific) connected in-line to an Easy-nLC HPLC system (Thermo Scientific), as described (53). The raw data were processed with Protein Discoverer software (Thermo Fisher Scientific, Inc.). Database searches were performed using the in-house Mascot server (Matrix Science) against a database of *Synechocystis* 6803 proteins supplemented with sequences of common protein contaminants. The search criteria allowed for one miscleavage of trypsin, oxidation of methionine, acetylation of the protein N-termini and phosphorylation of S, T and Y residues.

### Determination of NAD^+^/NADH

All the cultures used for NAD^+^/NADH determination experiment were grown autotrophically and mixotrophically in 250 ml BG-11 medium. 5 ml to 10 ml cells, equivalent to about 10^9^ cells/ml (10 ml cultures of OD_750_ of 1) were sampled for the measurements. The cells were centrifuged at 3,500 x g -9 °C for 10 min and the pellets were washed with 1 ml 20 mM cold PBS (20 mM KH2PO4, 20 mM K2HPO4, and 150 mM NaCl). The suspension was transferred to a 2 ml reaction cup and was centrifuged at 12,000 x g for 1 min at −9 °C. For all further steps the NAD+/NADH Quantification Colorimetric Kit (Biovision, CA, USA ) was used. The pellet was resuspended in 50 μl extraction buffer and precooled glass beads (∅=0.18 mm) were added to about 1 mm to the surface of the liquid. The mixture was vortexed 4 times 1 min in the cold room (4 °C) and intermittently chilled on the ice for 1 min. 150 μl extraction buffer was added again and the mixture was centrifuged at 3,500 xg for 10 min at −9 °C. The liquid phase was transferred as much as possible into a new reaction cup and centrifuged at maximum speed for 30 min at −9 °C. All further steps were conducted as described by the manufacturer. Finally, the samples were incubated for 1 to 4 hours in 96 well plates before measuring absorbance at 450 nm by TECAN GENios (TECAN Group Ltd., Austria) along with a NADH standard curve.

### Immunoblots

Whole cell extracts were applied on 10 % SDS-polyacrylamide gels according to the protocol of Laemmli (1970). After separation of the proteins the gel was semi-dry blotted on a nitrocellulose membrane (Roti-NC, 0.2 μm Transfer membrane for protein analyses, Carl Roth, Karlsruhe, Germany). For blocking the membrane was incubated in 2.5 % BSA in TBS (20 mM Tris pH 7.5, 150 mM NaCl). For the detection of phosphorylated Serine residues a TBS solution with a 1:100 dilution of the Anti-Phospho-Serin Antibody (Qiagen, Hilden, Germany) was used. For detection of the PdhA subunit an antibody was raised against the peptide TKYRREVLKDDGYDQ (Fig. S7) in rabbits by Agrisera (Umeå, Sweden). On the membranes 1:1000 dilution of this antibody in TBS was used. As secondary antibody either an anti-mouse IgG-HRP conjugate or an anti-rabbit IgG-HRP conjugate was used at 1:10,000 dilution. For detection the membrane was immersed in a 1:1 mixture of solution A (100 mM Tris/HCl pH = 8.5, 0.4 mM p-coumaric acid, 2.5 μM luminol) and solution B (100 mM Tris/HCl pH = 8.5, 100 mM H_2_O_2_) for 1 min and subsequently exposed to an X-ray film (Thermo Scientific CL-XPosure Film, Life Technologies GmbH, Germany).

### Antibody against the PdhA subunit of the PDH complex

In order to raise an antibody against the alpha subunit of PdhA from the PDH complex, the peptide TKYRREVLKDDGYDQ (Fig. S7) was used for immunization of rabbits (Agrisera, Umeå, Sweden).

### Purification and activity measurement of dihydrolipoyl dehydrogenase (E3 subunit, SynLPD)

The recombinant His-tagged SynLPD (Slr1096) was generated and purified essentially as described previously (54). Prior activity measurements, the elution fractions were desalted through PD10 columns (GE healthcare, Solingen, Germany). The protein concentration was determined according to Bradford (55). SynLPD activity was determined in the forward direction. DL-dihydrolipoic acid served as the substrate at a final concentration of 3 mM. SynLPD activity was followed as reduction of NAD^+^ (included in varying concentrations, 0.1, 0.2, 0.3, 0.4, 0.5, 1, 2, 3, 4 and 5 mM) at 340 nm. The *K_i_* constant was estimated in the presence of four NADH concentrations (0, 0.1, 0.15 and 0.2 mM) as well as NADPH (0.1 mM) as control. Specific enzyme activity is expressed in µmol NADH per min^−1^ mg protein^−1^ at 25°C. Mean values and standard deviations were calculated from at least three technical replicates for all substrate/co-substrate combinations. All chemicals were purchased from Merck (Darmstadt, Germany).

### Purification of pyruvate:ferredoxin oxidoreductase (PFOR)

For the purification of PFOR from *Synechocystis* sp. PCC 6803 (Fig. S9, S10), three 6-L autotrophic cultures of the PFOR overexpression strain (PFOR:oe) were grown to an OD_750_ of about 1. Cells were harvested by centrifugation at 4.000 rpm in a JLA-8.1000 rotor for 20 min at 4°C. Initially, His-PFOR over-expression in the 6-L cultures was assessed by SDS PAGE analysis followed by immunoblotting with a His-tag specific antibody (GenScript; Fig S7). A specific band could be detected in the over-expression mutant, confirming expression and stable accumulation of the over-expressed and N-terminally His-tagged PFOR protein. For large-scale purification cells were resuspended in lysis buffer (50 mM NaPO_4_ pH=7.0; 250 mM NaCl; 1 tablet complete protease inhibitor EDTA free (Roche, Basel, Switzerland) per 50 mL) and broken by passing them through a French Press cell at 1250 p.s.i. twice. Unbroken cells and membranes were pelleted in a Beckman ultracentrifuge using a 70 Ti rotor at 35.000 rpm for 45 min at 4°C. The decanted soluble extract was adjusted to a volume of 90 mL with lysis buffer and incubated with 10 mL TALON cobalt resin (Takara, Shiga, Japan) for 1 h at 4°C. The resin was then washed extensively with 200 mL lysis buffer and subsequently with 100 mL lysis buffer containing 5 mM imidazole. Bound proteins were eluted with 20 mL elution buffer (50 mM NaPO_4_ pH=7.0; 250 mM NaCl; 500 mM imidazole). The protein was concentrated overnight to a volume of 2 mL in a Vivaspin 20 Ultrafiltration Unit (5 kDa MWCO)(Merck, Darmstadt, Germany) and then loaded onto a HiLoad^TM^ 26/60 Superdex TM 75 prep grade (GE Healthcare, Chicago, IL, USA) using 25 mM NaPO_4_, pH=7.0; 50 mM NaCl; 5% (v/v) glycerol as the running buffer. The run was monitored at 280 nm and fractions were collected (Fig. 8A).

### Acitivity measurement of pyruvate:ferredoxin oxidoreductase (PFOR)

The specific activity of the pyruvate:ferredoxin oxidoreductase was measured essentially as described (11). The activity assay contained in 1 ml 100 mM Tris-HCl (pH 8), 0.5 mM Coenzyme A, 10 mM pyruvate, 5 mM thiamine pyrophosphate, 40 mM glucose, 40 U glucose oxidase, 50 U catalase, and 10 mM methy viologen. Reduction of methylviologen was followed at 604 nm and an extinction coefficient of 13.6 mM^−1^ cm^−1^ was used. The reaction was started by adding 8.9 x 10^− 5^ M isolated PFOR.

We also tested ferredoxin reduction by the PFOR by a mixture containing the same substances as above except methyl viologen. To this mixture 1.6 mM ferredoxin 1 and 1.3 mM ferredoxin:NADP^+^ reductase and 1 mM NADP^+^ were added. In this case the reduction of NADP^+^ was followed at 340 nm. The same mixture without glucose, glucose oxidase and catalase were used to test if the enzyme also works in the presence of oxygen.

### In-vivo electron flow through photosystem I

The electron flux through photosystem I was measured by the Dual-KLAS/NIR (Walz GmbH, Effeltrich, Germany) by a newly developed method (36). In brief, cell suspensions were adjusted to 20 µg/mL chlorophyll and 20 µM DCMU was added. Electron flow through PSI was determined by dark-interval relaxation kinetics (DIRK) measurements at a light intensity of 168 µE/m^2^/s in the absence and presence of 10 mM glucose.

### Determination of reaction Gibbs energies

Δ_r_G’^m^ for the reaction catalyzed by the PDH complex and by PFOR were calculated using eQuilibrator (http://eauilibrator.weizmann.ac.il/) according to (44). CO_2_ (total) was considered as hydrated and dehydrated forms of CO_2_ are considered to be in equilibrium in biochemical reactions. Ionic strength of 0.2M, pH of 7 and metabolite concentrations of 1 mM were assumed. In order to determine the redox potential of pyruvate we used the reactions Gibbs energy of -39 kJ/mol for the PDH complex and -23 kJ/mol for PFOR. Assuming a redox potential of -320 mV for NAD(P)H and -400 mV for ferredoxin the potential of pyruvate was determined according to ΔG = -nFΔE to -520 mV.

## Acknowledgments

Technical assistance of Klaudia Michl during the growth experiments of serine/threonine protein kinase (spk) deletion mutants is acknowledged.

This study was supported by grants from the China Scholarship Council (CSC) (Grant # 201406320187), FAZIT-Stiftung, Deutsche Bundesstiftung Umwelt, German Ministry of Science and Education (BMBF FP309), and the German Science Foundation (DFG Gu1522/2-1, HA2002/23-1 and FOR2816).

## Supplementary Information

**Figure S1:**
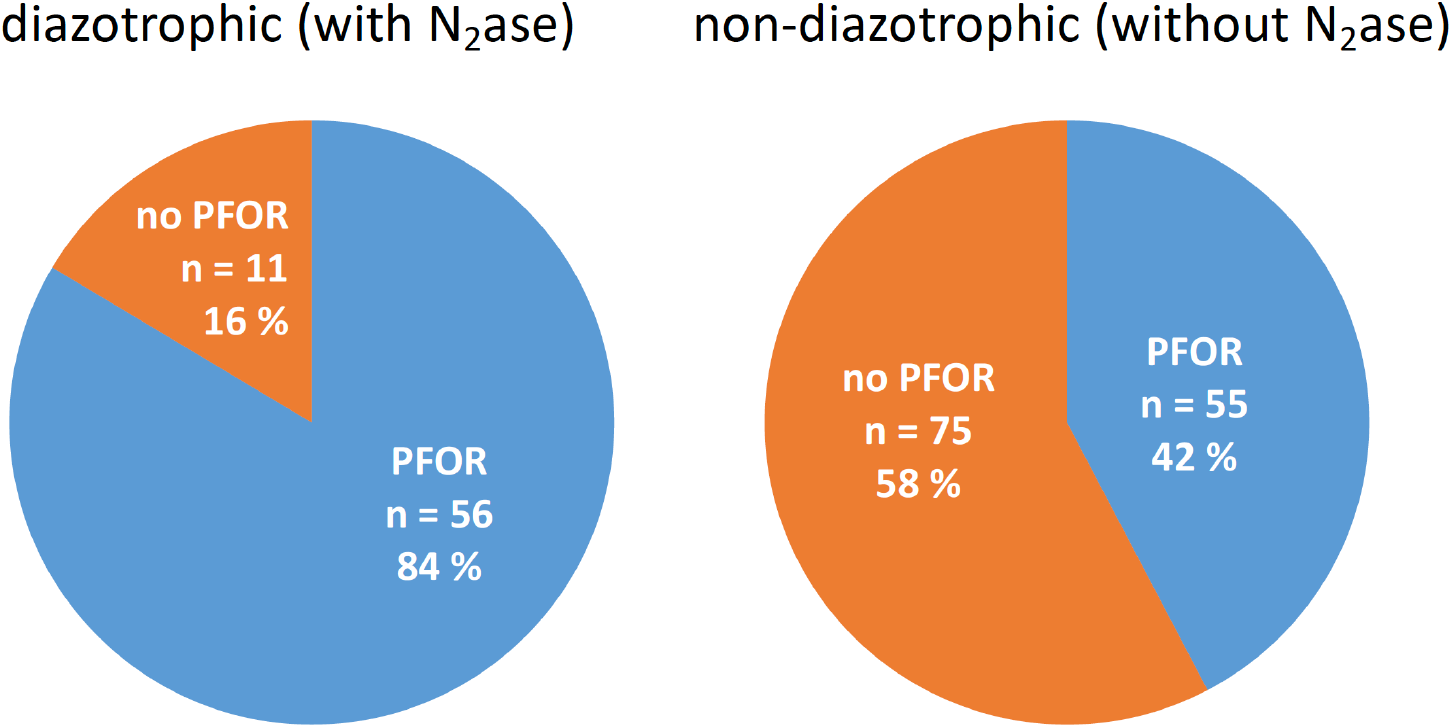
Bioinformatic analyses concerning the distribution of PDH complex and PFOR in diazotrophic and non-diazotrophic cyanobacteria. All shown genomes possess a PDH complex.

All completely sequenced cyanobacterial genomes were analyzed via tblastn for the presence of the PDH complex and PFOR. For this, in order to exclude symbionts, cyanobacterial genomes were in a first step searched for the *psbD* gene (PSII subunit). We used the *psbD* gene (*sll0849*) of *Synechcoystis* as bait. Only genomes containing *psaD* were used for all further analysis. 197 genomes remained and were searched by tblastn using the *pdhA* subunit (*slr1934*) from the PDH complex from *Synechcoystis* as bait. The largest expect value was 2×10^−136^. *pdhA* was found in all genomes analyzed. 67 of these genomes contain *nifD* (highest e-value 4×10^−104^) and *nifK* (highest e-value 1×10^−73^), the two subunits of the nitrogenase for N_2_-fixation and a diazotrophic lifestyle. Diazotrophic and non-diazotrophic cyanobacteria were searched for the presence of PFOR by using *sll0741* from *Synechcoystis*. The highest e-value in this case was 0.

We found that all fully sequenced diazotrophic and non-diazotrophic cyanobacteria with PSII contain genes coding for a PDH complex and that 56 % thereof possess a PFOR as well. If we subtract from this group all diazotrophic cyanobacteria that contain a nitrogenase and might therefore utilize PFOR in the process of nitrogen fixation, 130 non-diazotrophic cyanobacteria remain. Within the group of non-diazotrophic cyanobacteria 42% possess a PFOR in addition to the PDH complex. This clearly shows that the property of holding both a PDH complex and a PFOR in cyanobacteria that live predominantly under oxic conditions is truly widespread. The analysis furthermore confirms our observation, that the PDH complex is preferred over the utilization of PFOR in cyanobacteria.

**Figure S2:**
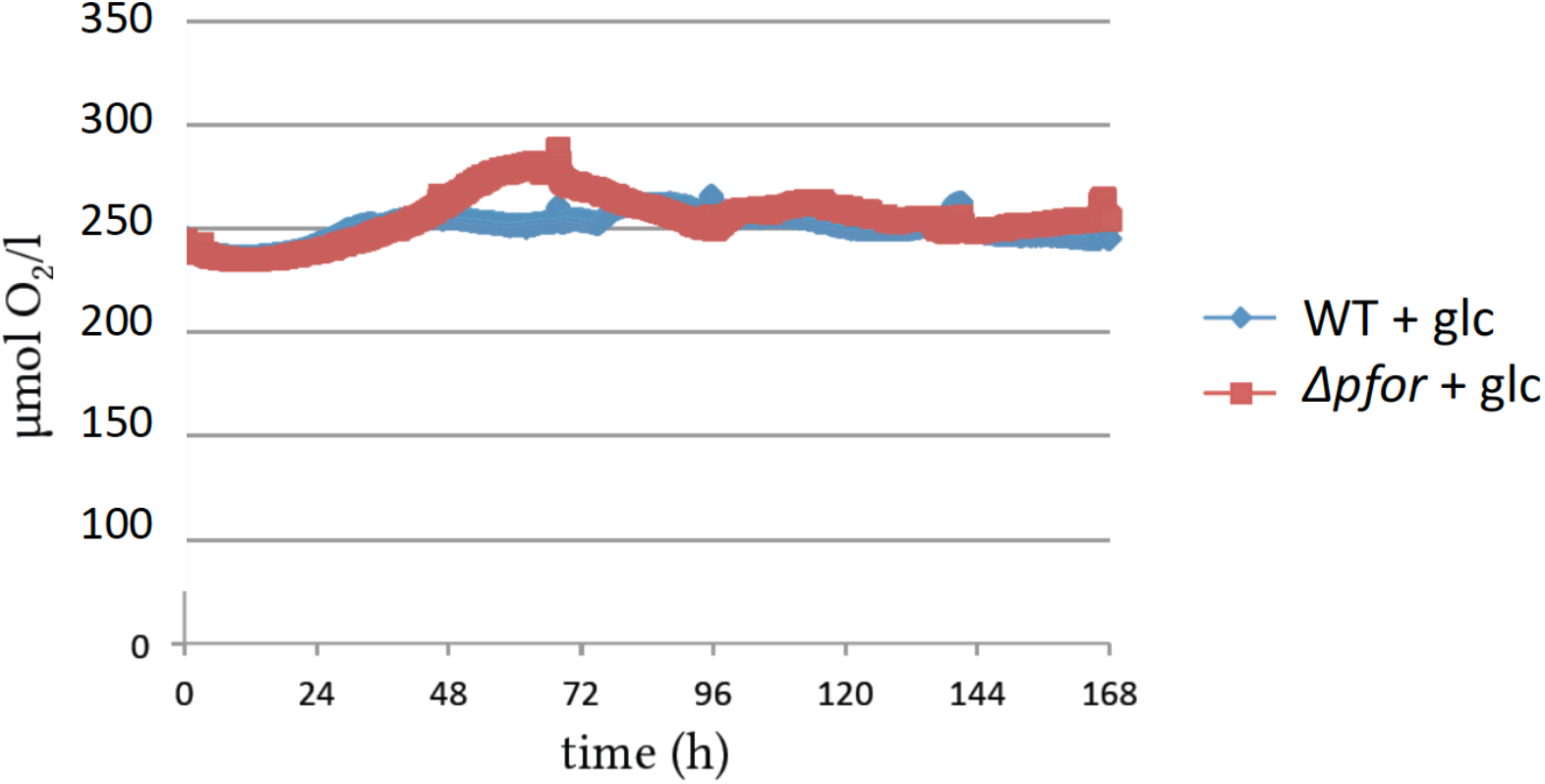
Oxygen concentrations in photomixotrophic cultures of wild type (WT) and Δ*pfor* were close to oxygen saturation throughout the growth experiments. Original traces are shown.

**Figure S3:**
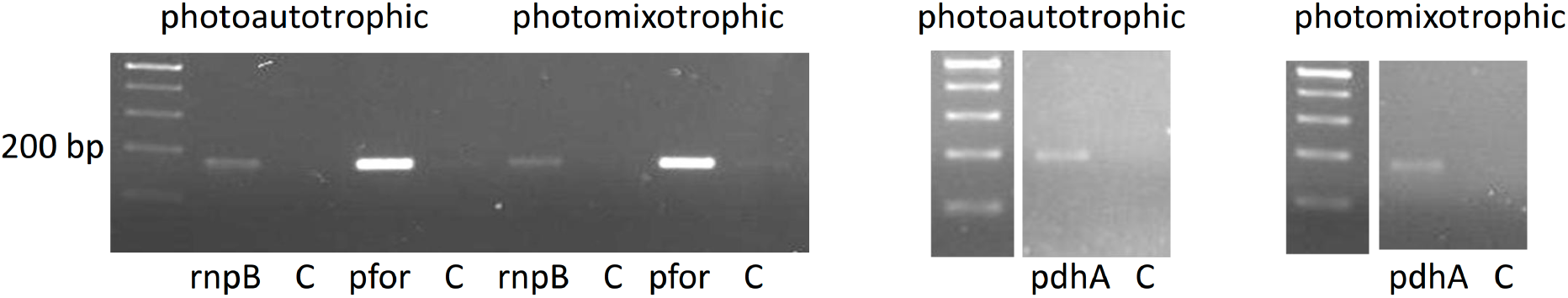
RT-PCR showing that *pfor* and *pdhA* are transcribed under photoautotrophic and photomixotrophic conditions in the wild type. Total RNA of wild type cells was reverse transcribed and subsequently subjected to PCRs with either primers specific for *rnpB*, *pfor* or *pdhA* (table S1). In the control reactions (C) reverse transcriptase was omitted.

**Figure S4:**
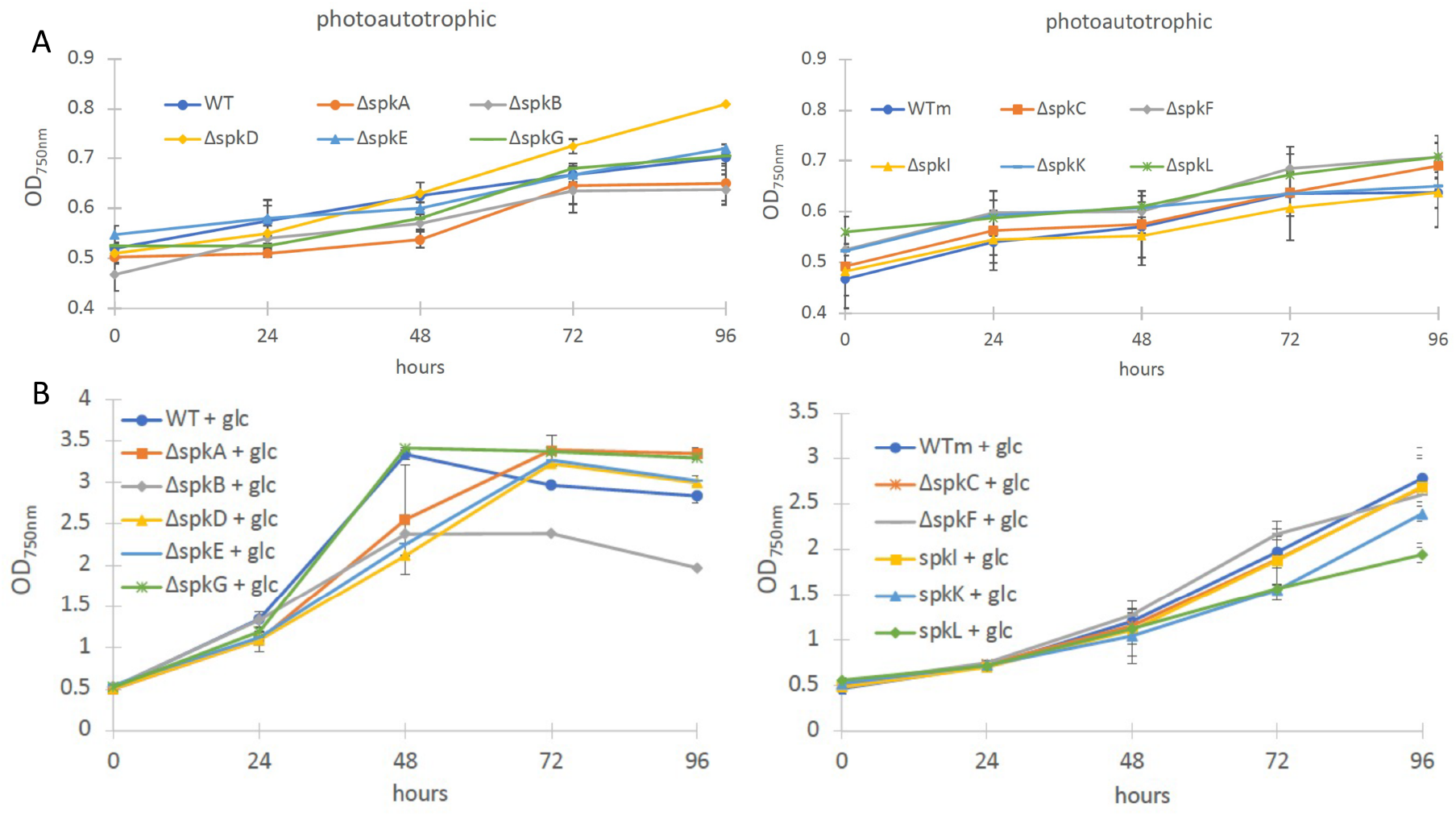
(A) Photoautotrophic and (B) photomixotrophic growth of serine/threonine protein kinase (spk) deletion mutants. The mutants were constructed in different *Synechocystis* WT backgrounds (table S2). *spkA*, *spkB*, *spkD*, *spkE* and *spkG* were deleted in the non-motile GT strain as all other mutants in this study with exception of *spkC*, *spkF*, *spkI*, *spkK*, and *spkL* that were deleted in the motile strain indicated as WTm. Shown are mean values ± SD from at least 3 replicates.

Photoautotrophic and photomoxotrophic growth of ten serine/threonine protein kinase (spk) deletion mutants was analyzed. They grew like the WT under photoautotrophic conditions (Fig. S4A) whereas the growth of Δ*spkB* and Δ*spkL* was affected under photomixotrophic conditions (Fig. S4B). This indicates that phosphorylation of enzymes is relevant for optimal photomixotrophic growth in *Synechocystis*.

**Figure S5:**
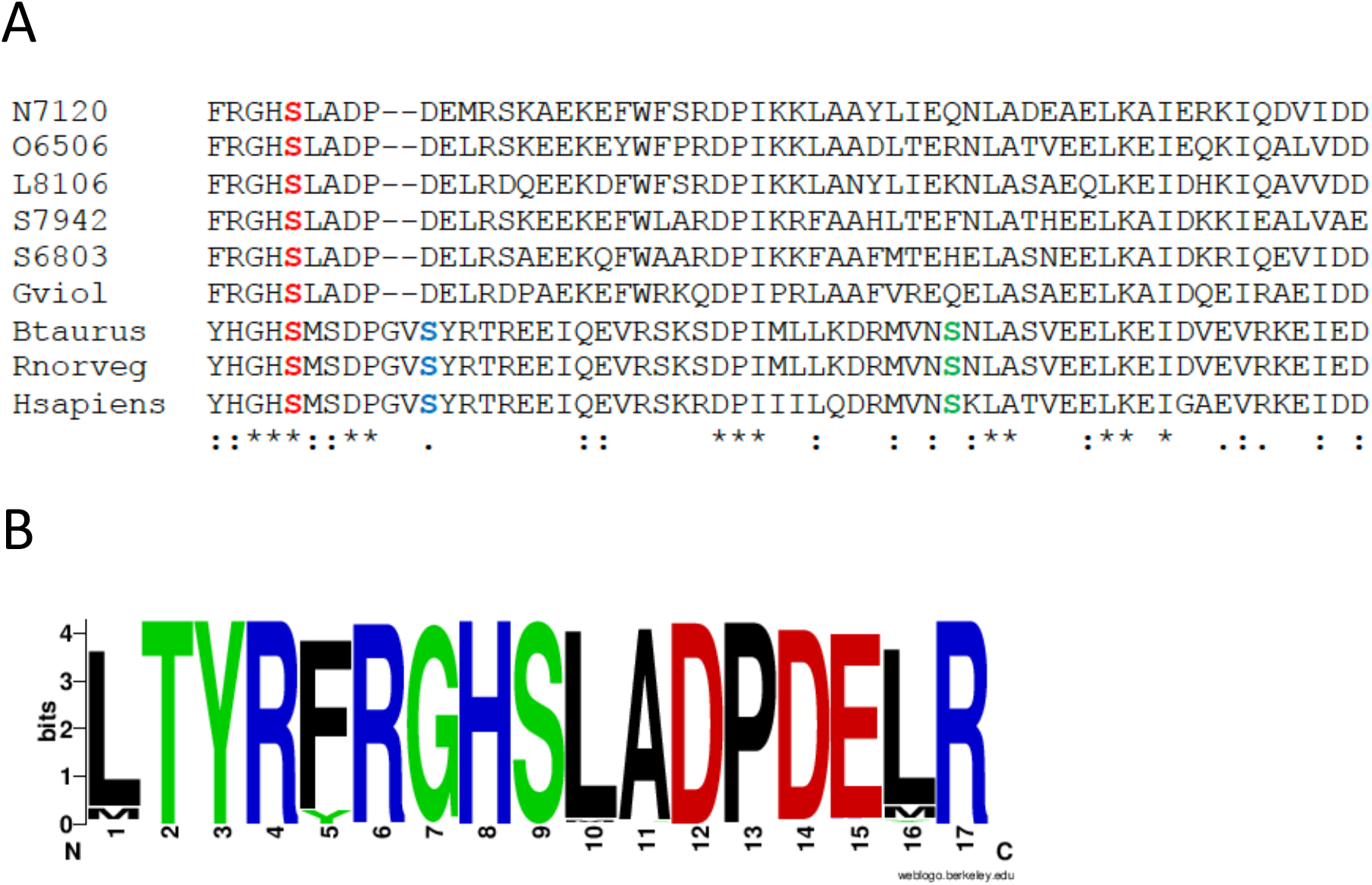
(A) Extract of the sequence alignment with the E1 subunits of the PDH complex. Shown are cyanobacterial sequences of *Nostoc* sp. PCC 7120 (N7120), *Oscillatoria* sp. PCC 6506 (O6506), *Lynbya* sp. PCC 8106 (L8106), *Synechococcus* sp. PCC 7942 (S7942), *Gloeobacter violaceus* (Gviol), and *Synechocystis* sp. PCC 6803 (S6803), and eukaryotic sequences of cattle (*Bos taurus* (Btaurus)), rat (*Rattus norvegicus* (Rnorveg)) and human (*Homo sapiens* (Hsapiens)). Serine residue 1 (**S**), is conserved in all PdhAs whereas residues 2 (**S**) and 3 (**S**), are only found in the eukaryotes. The first serine residue (**S**) was found in all 932 cyanobacterial PdhA sequences extracted from Genbank by a blast search as of January 11^th^ 2021 (Fig. S5). (B) Sequence logo of 17 amino acids around serine residue 1 from 932 cyanobacterial *pdhA* sequences showing that this region is highly conserved. The logo was generated via https://weblogo.berkeley.edu/logo.cgi.

**Figure S6:**
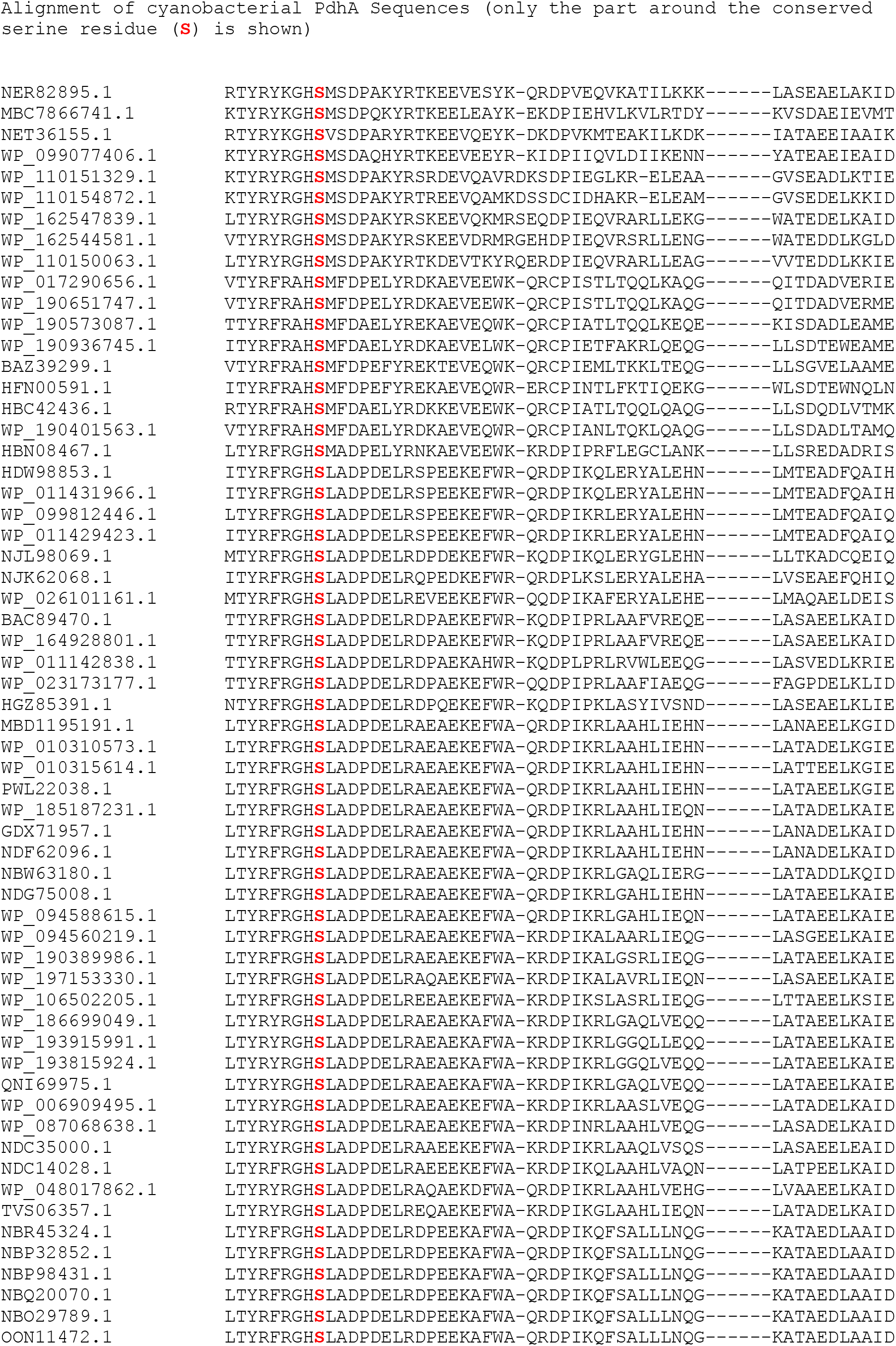
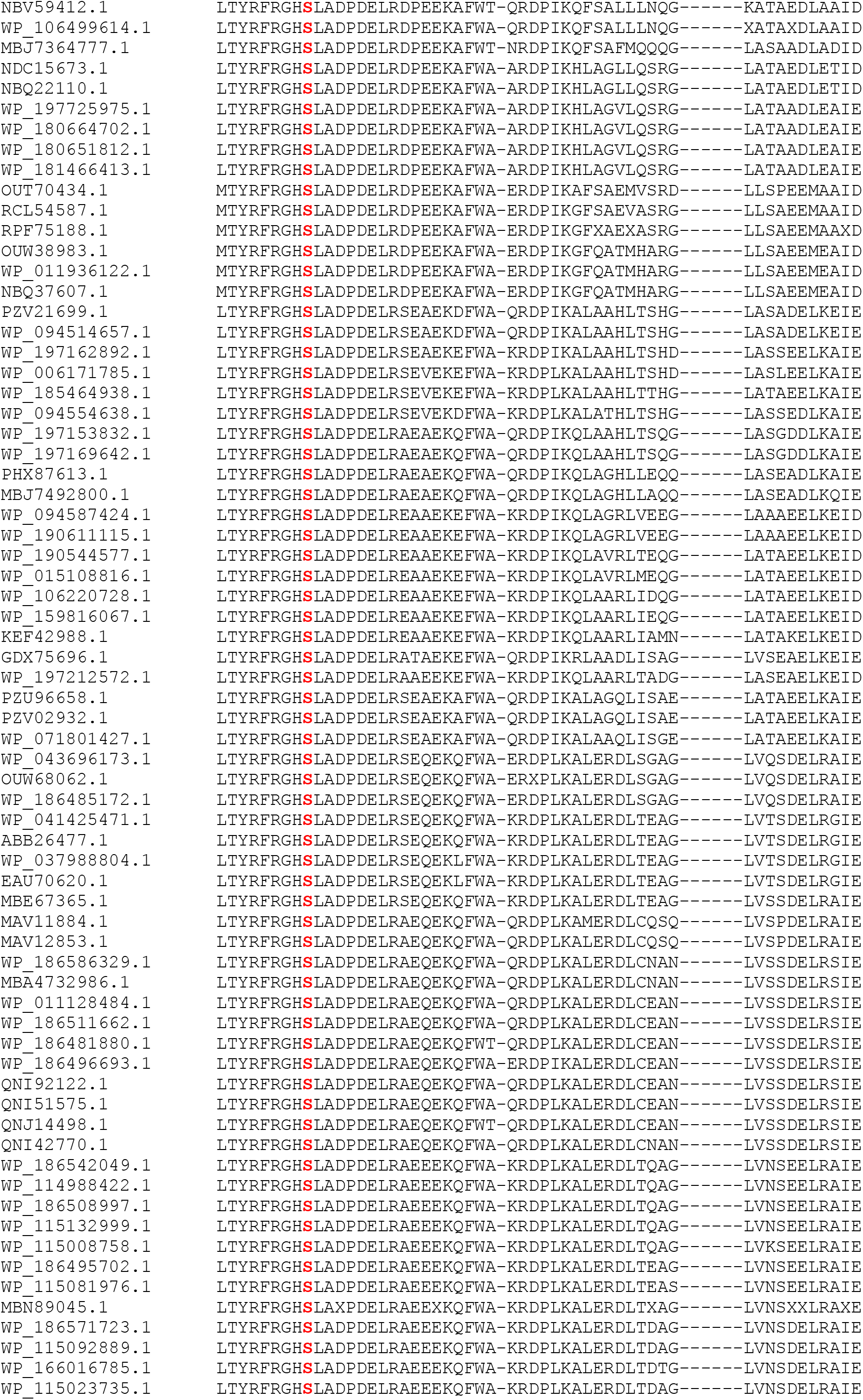
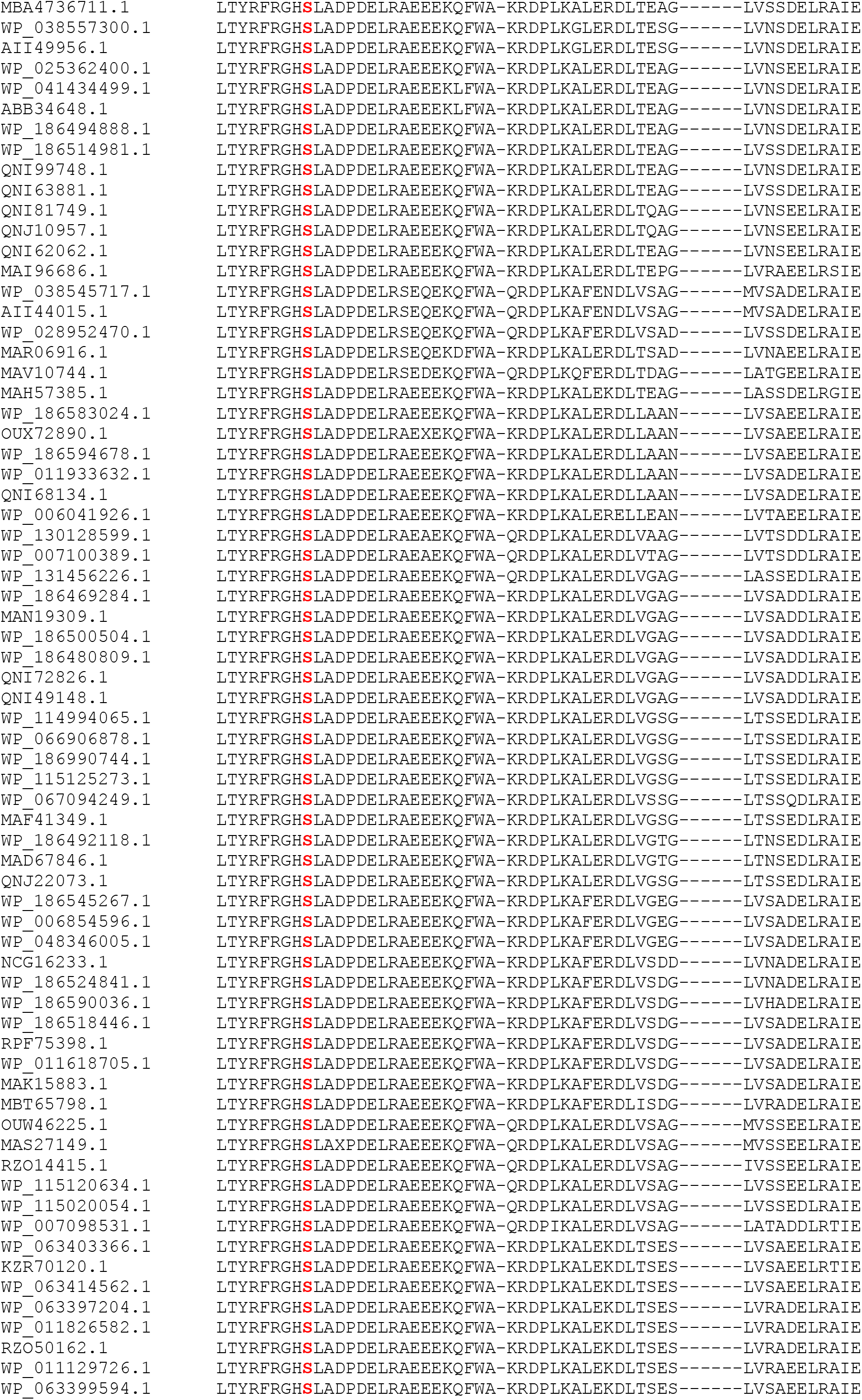
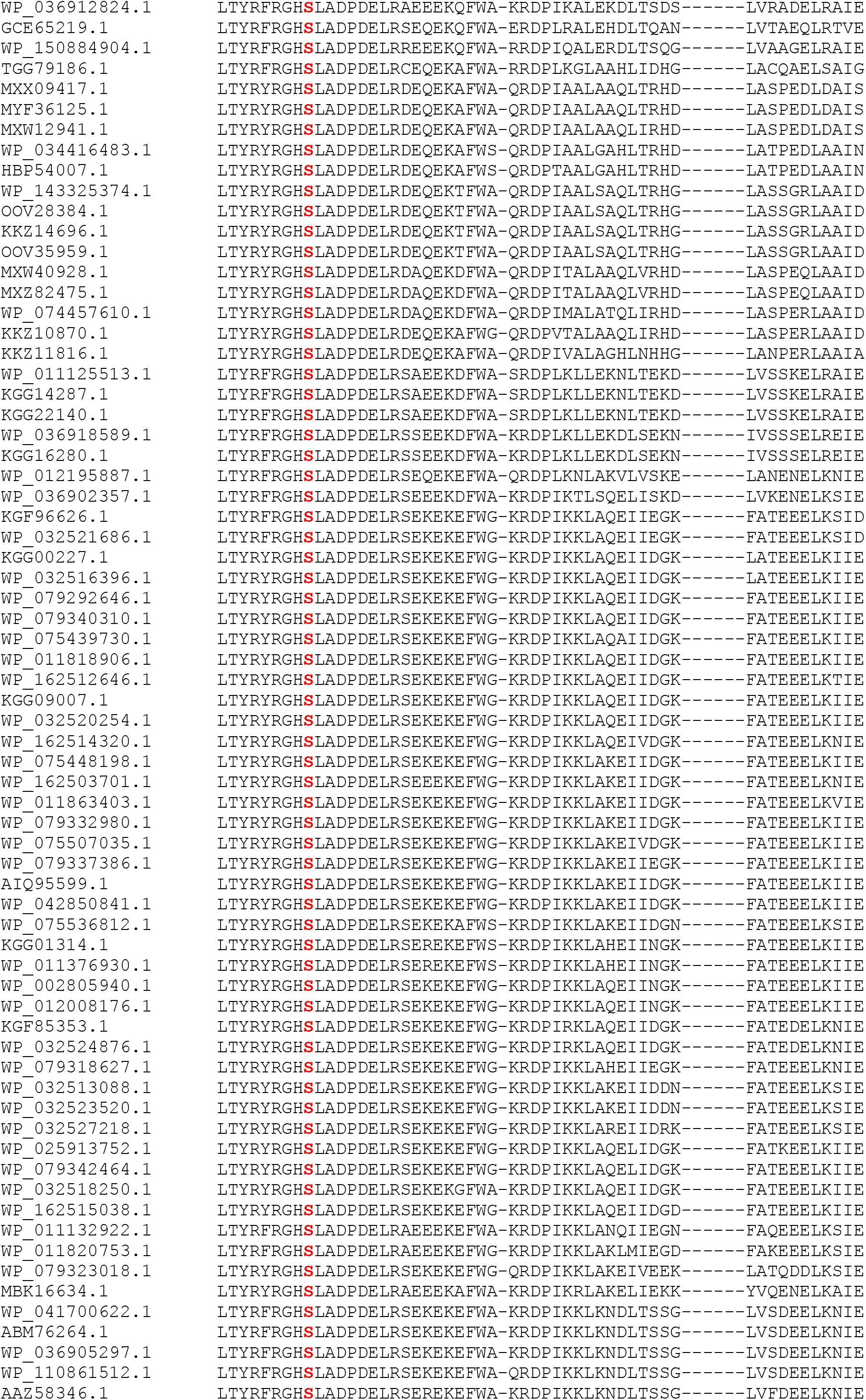
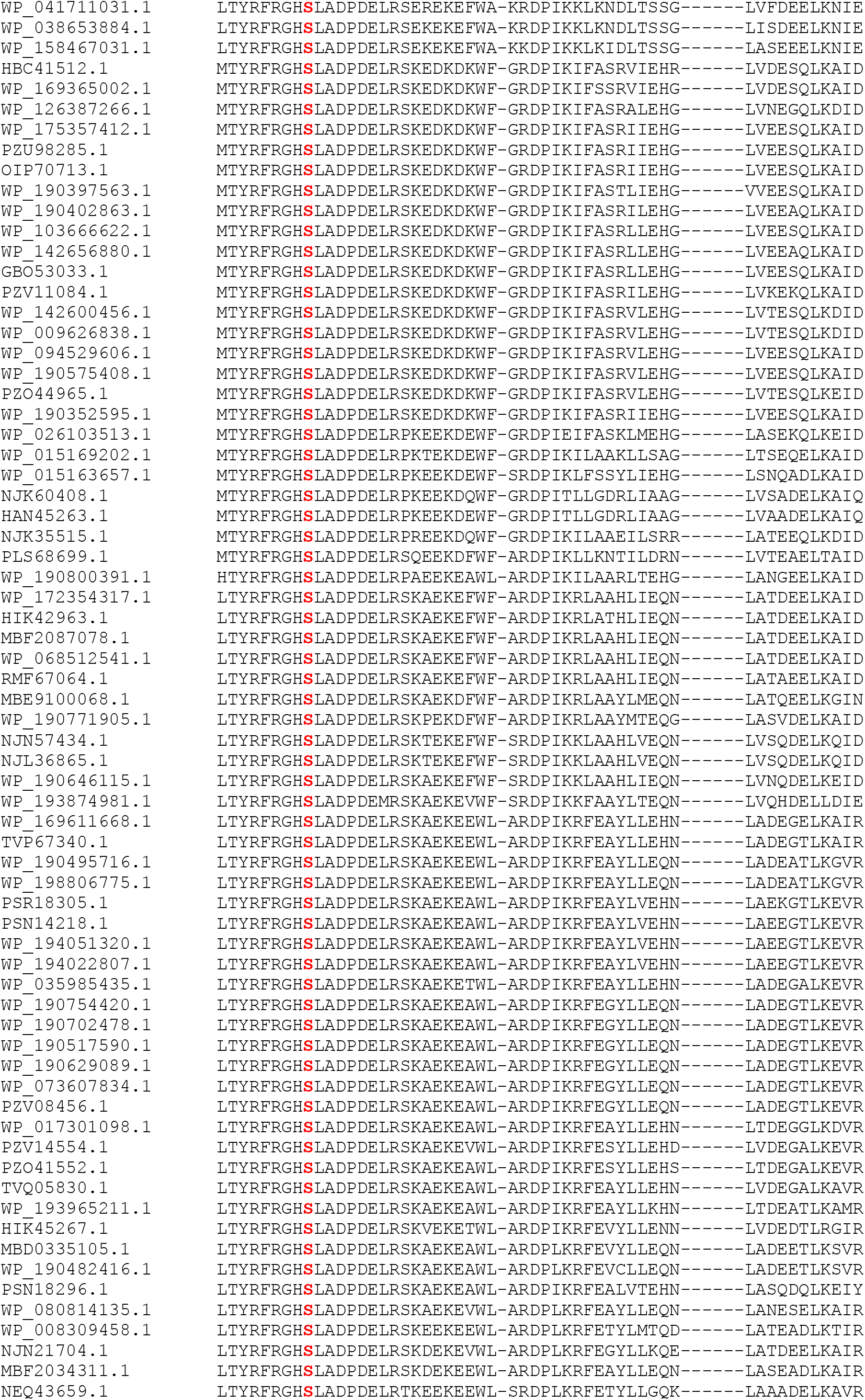
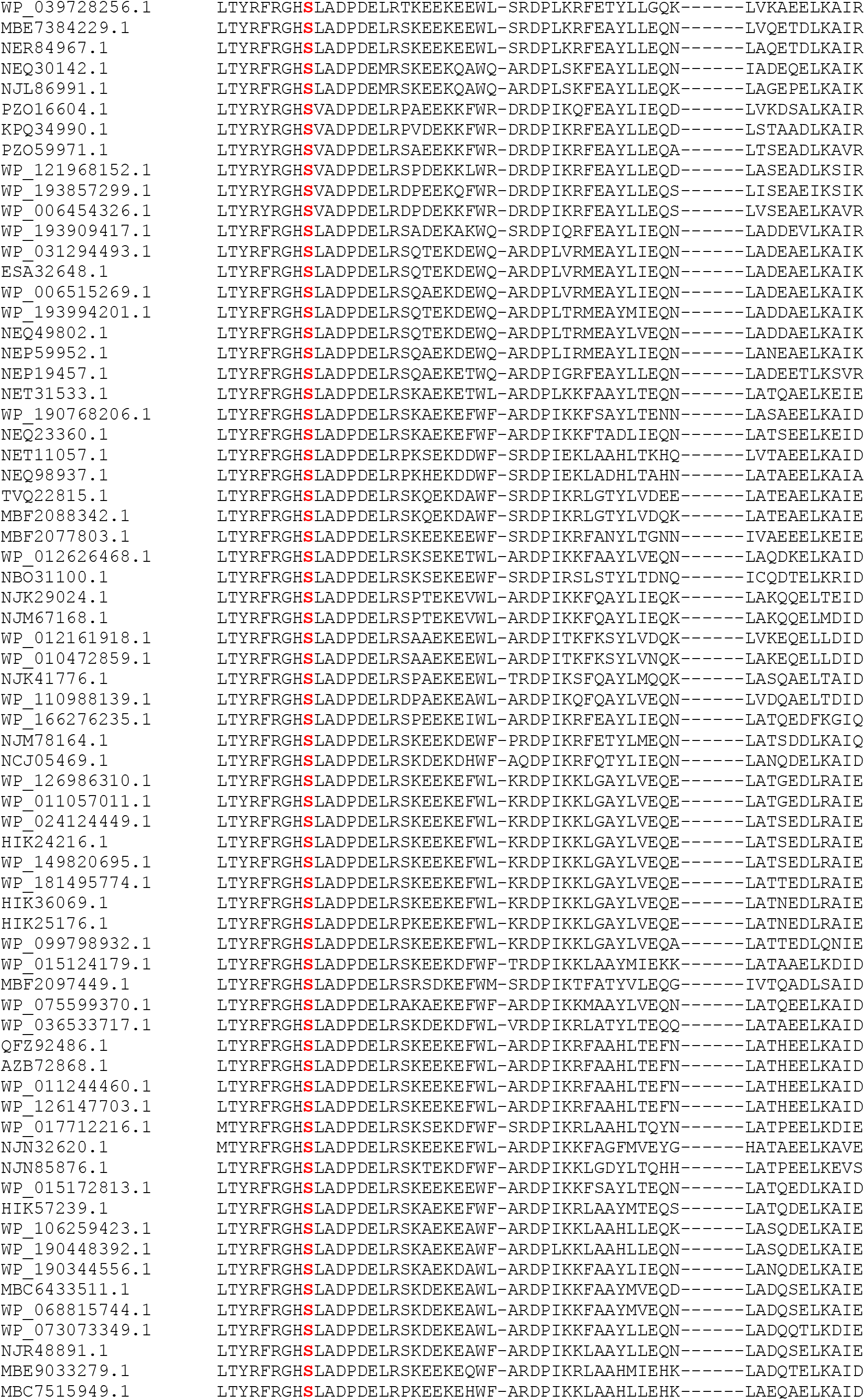
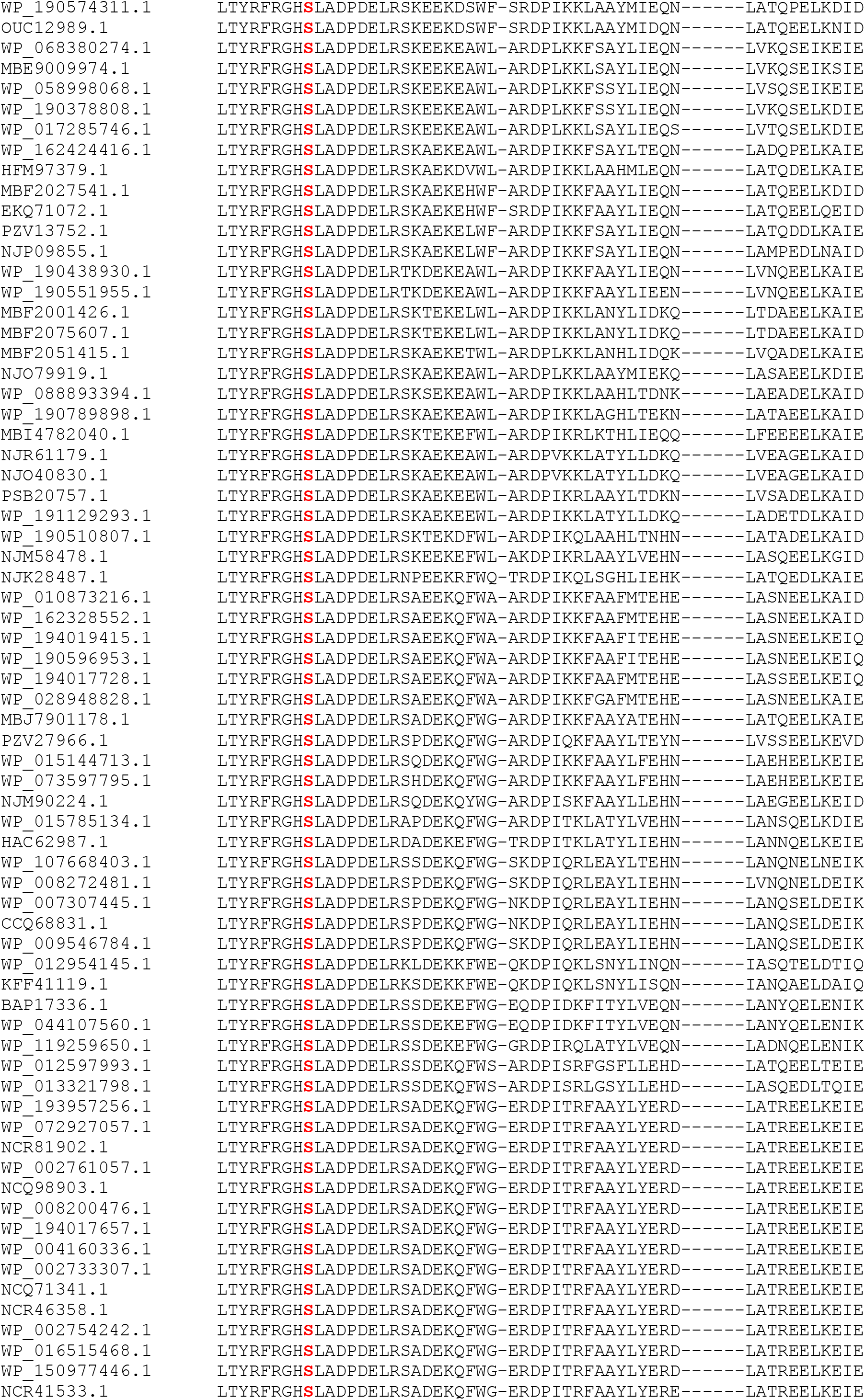
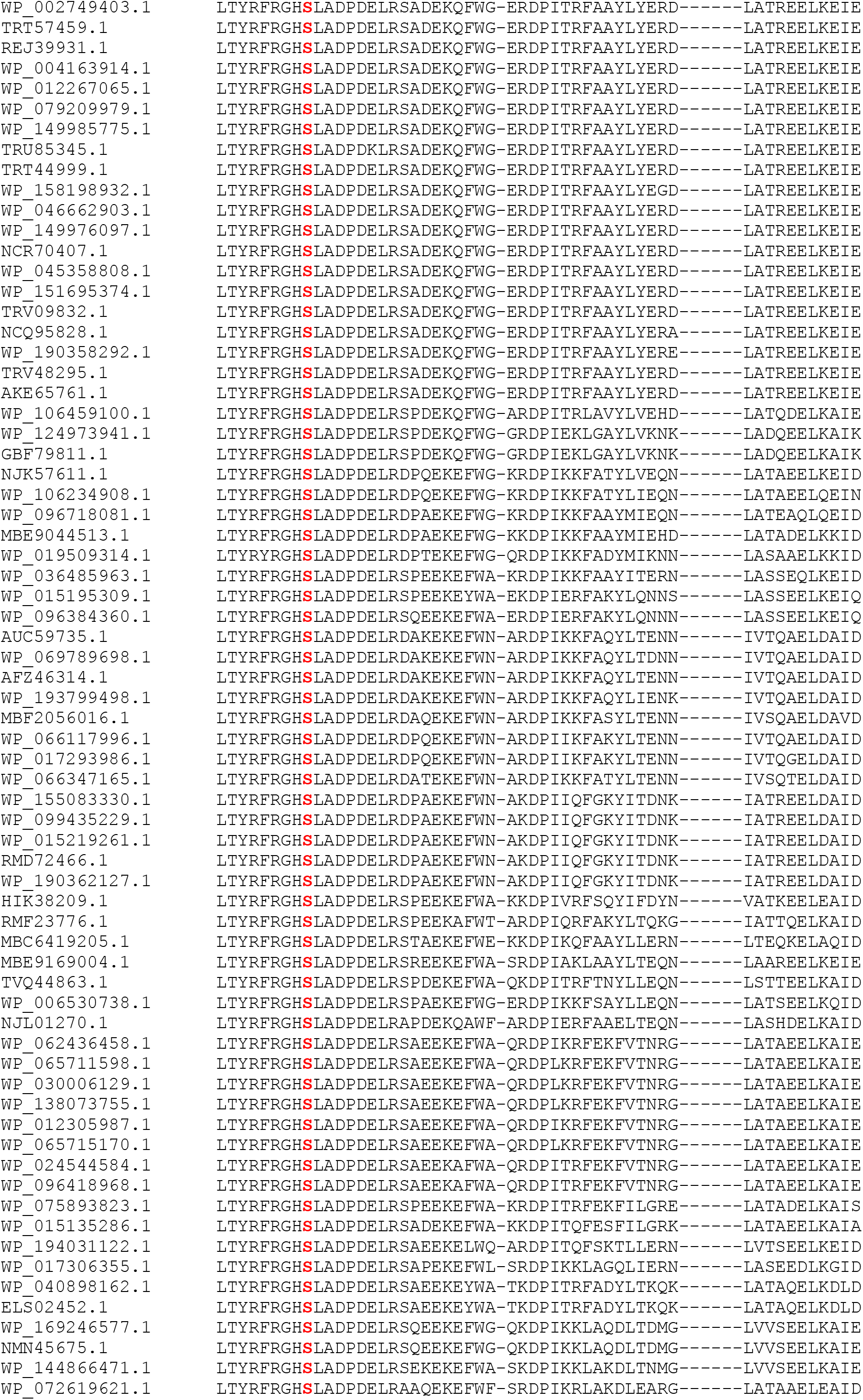
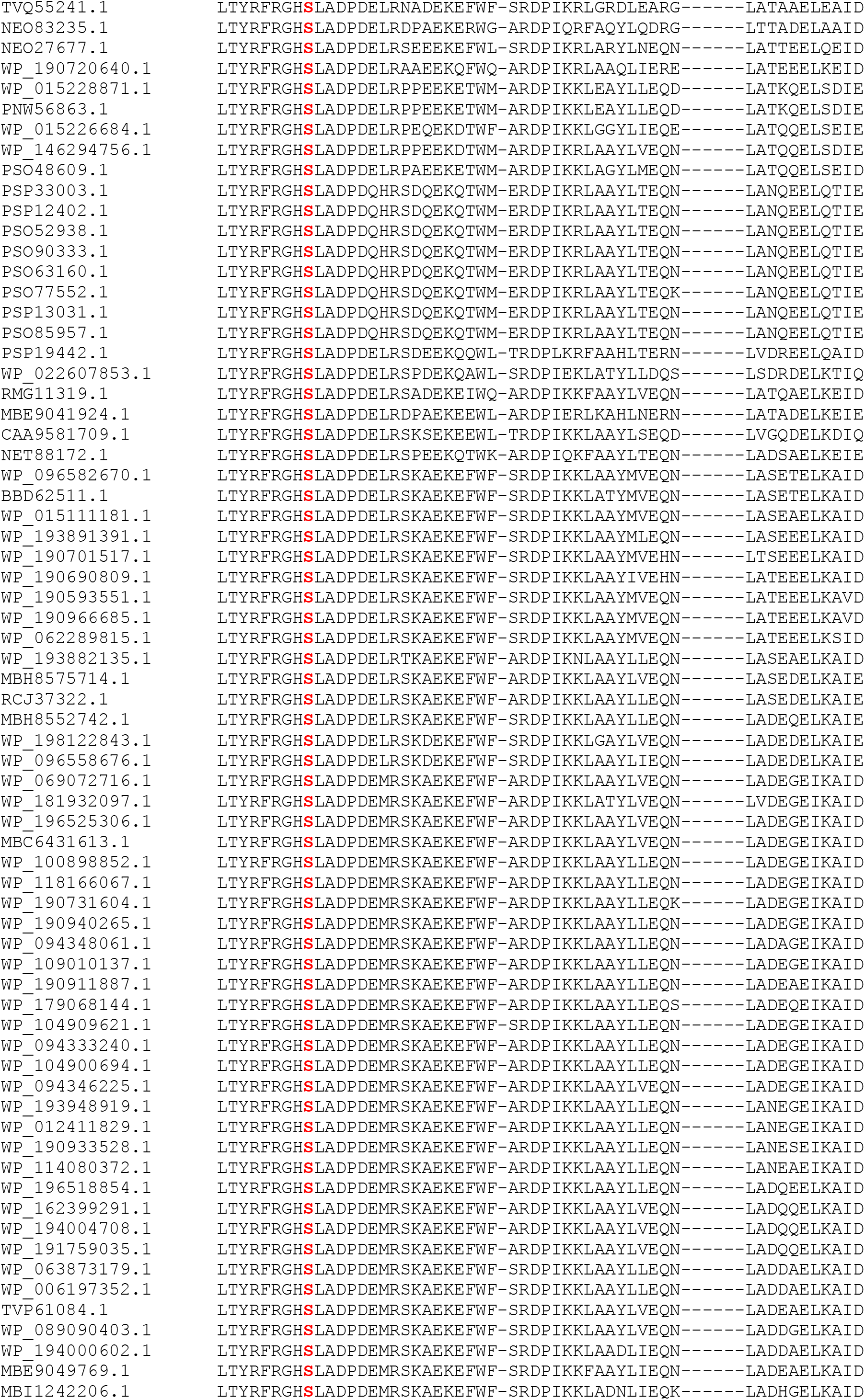
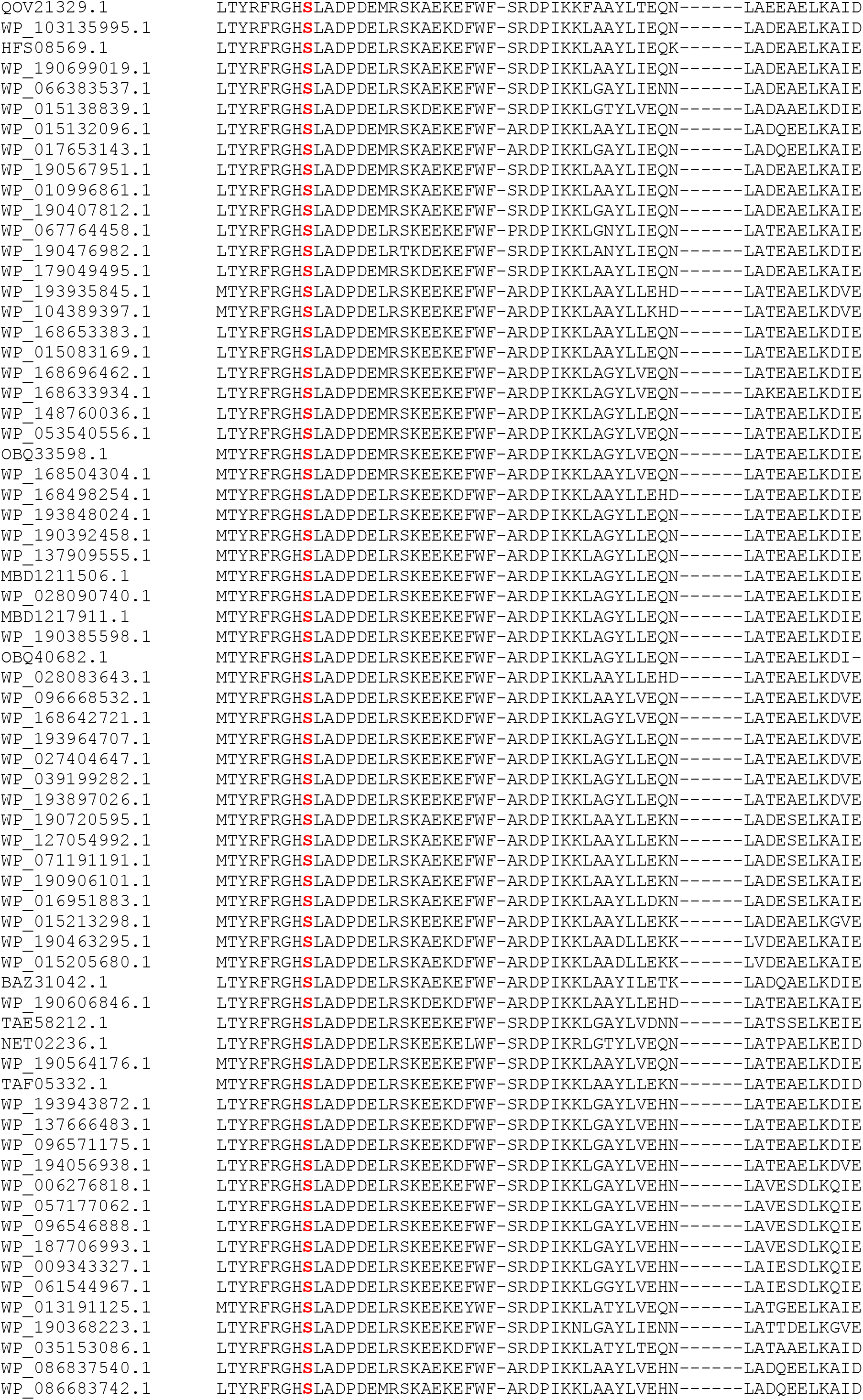
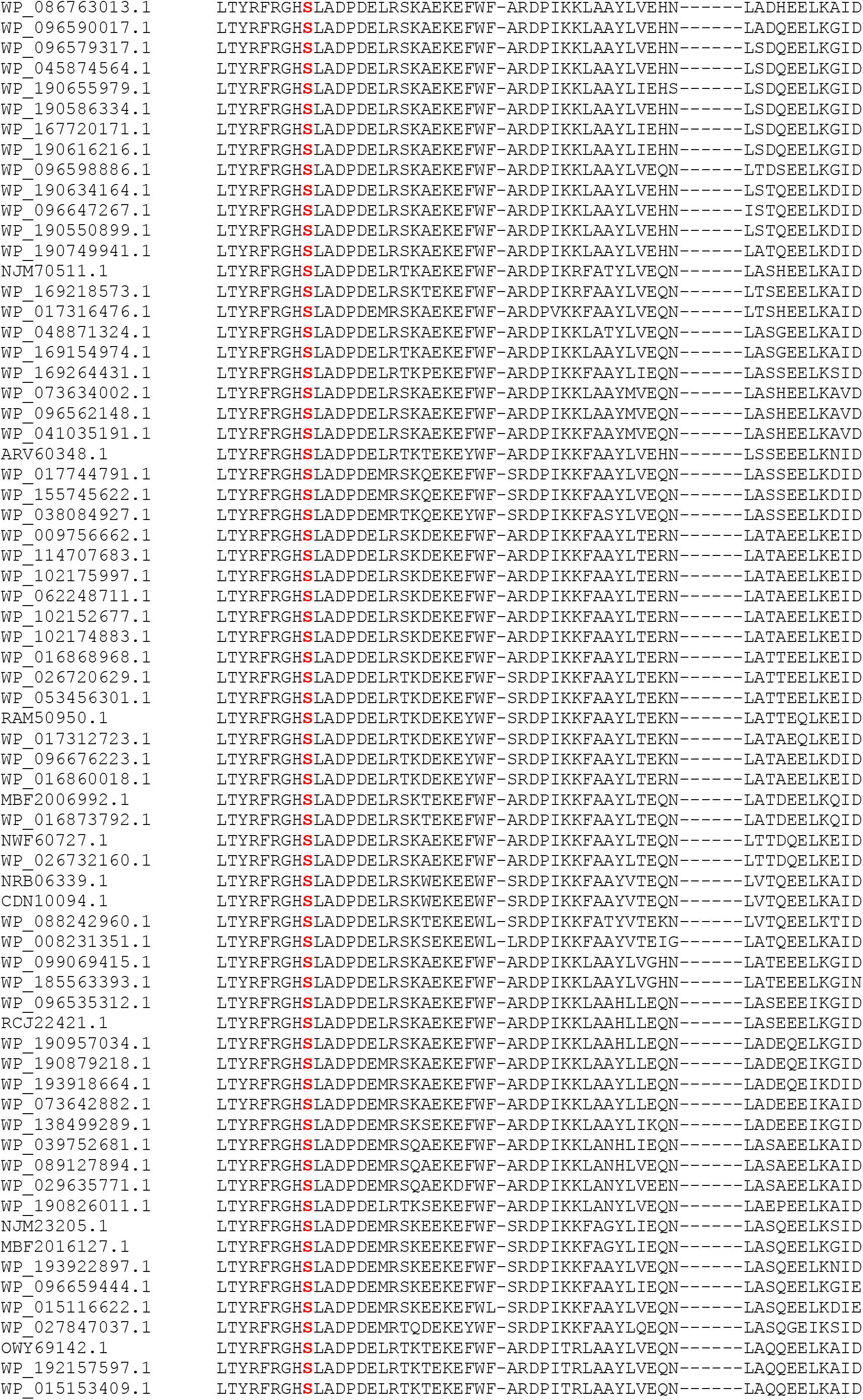
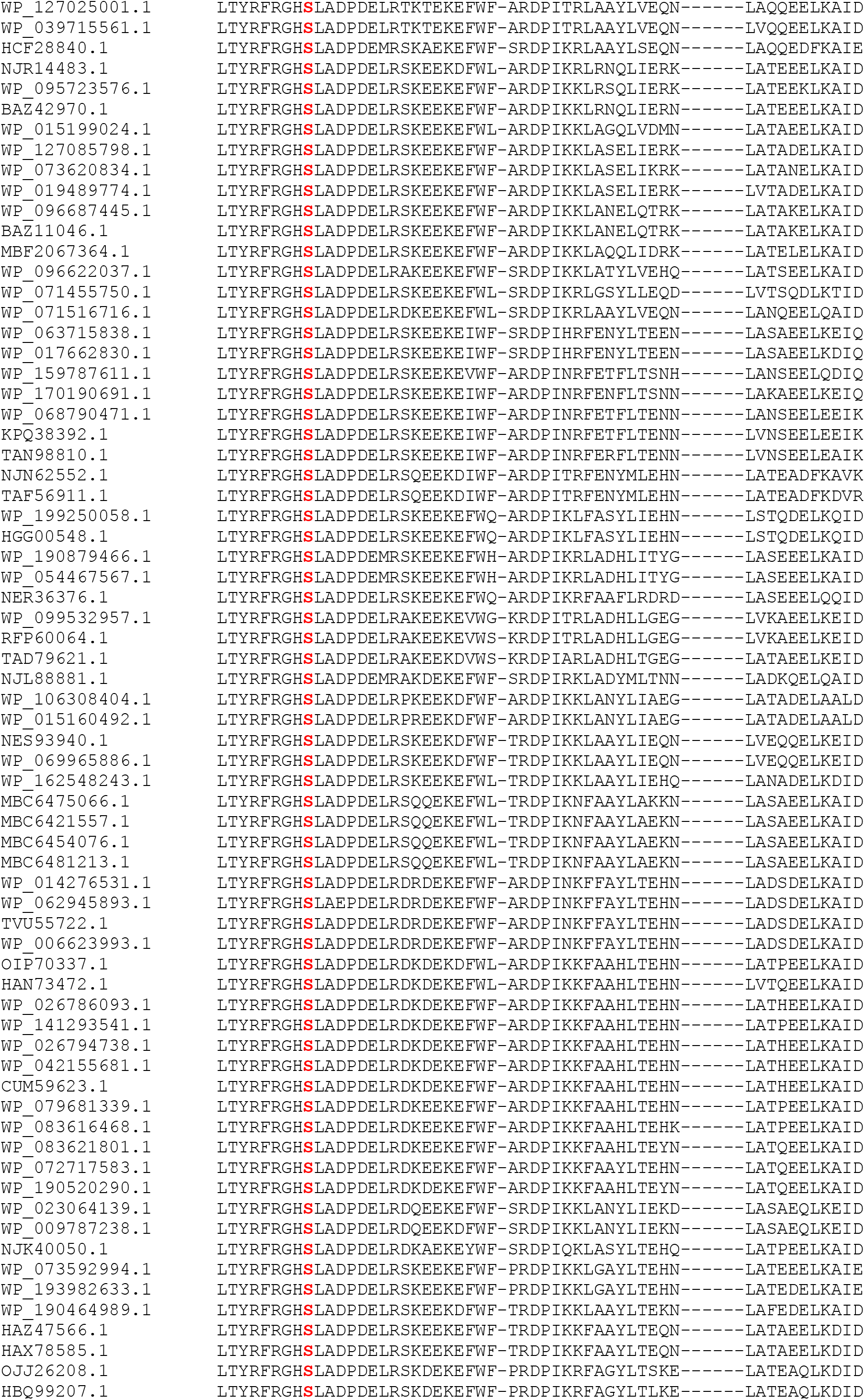
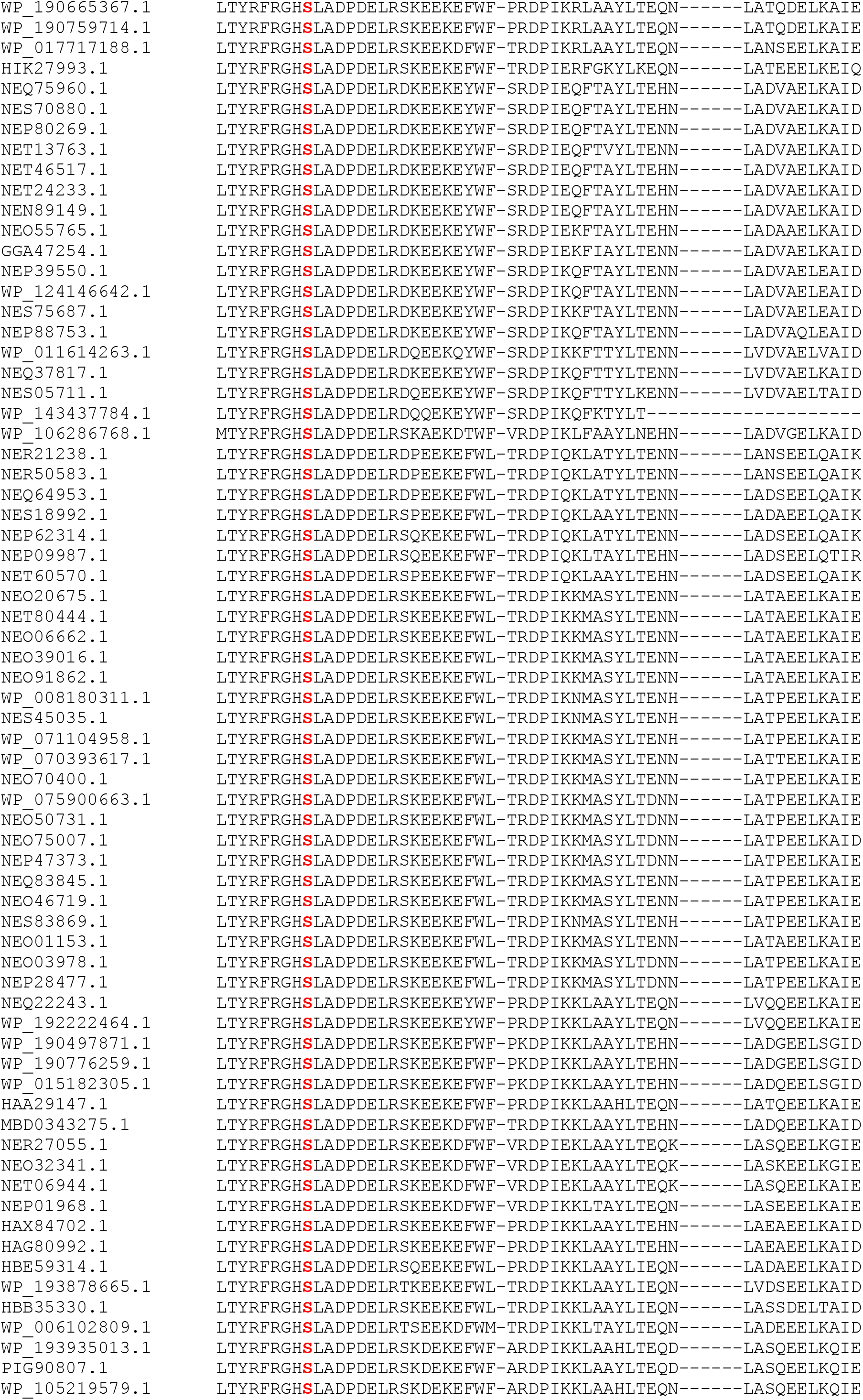
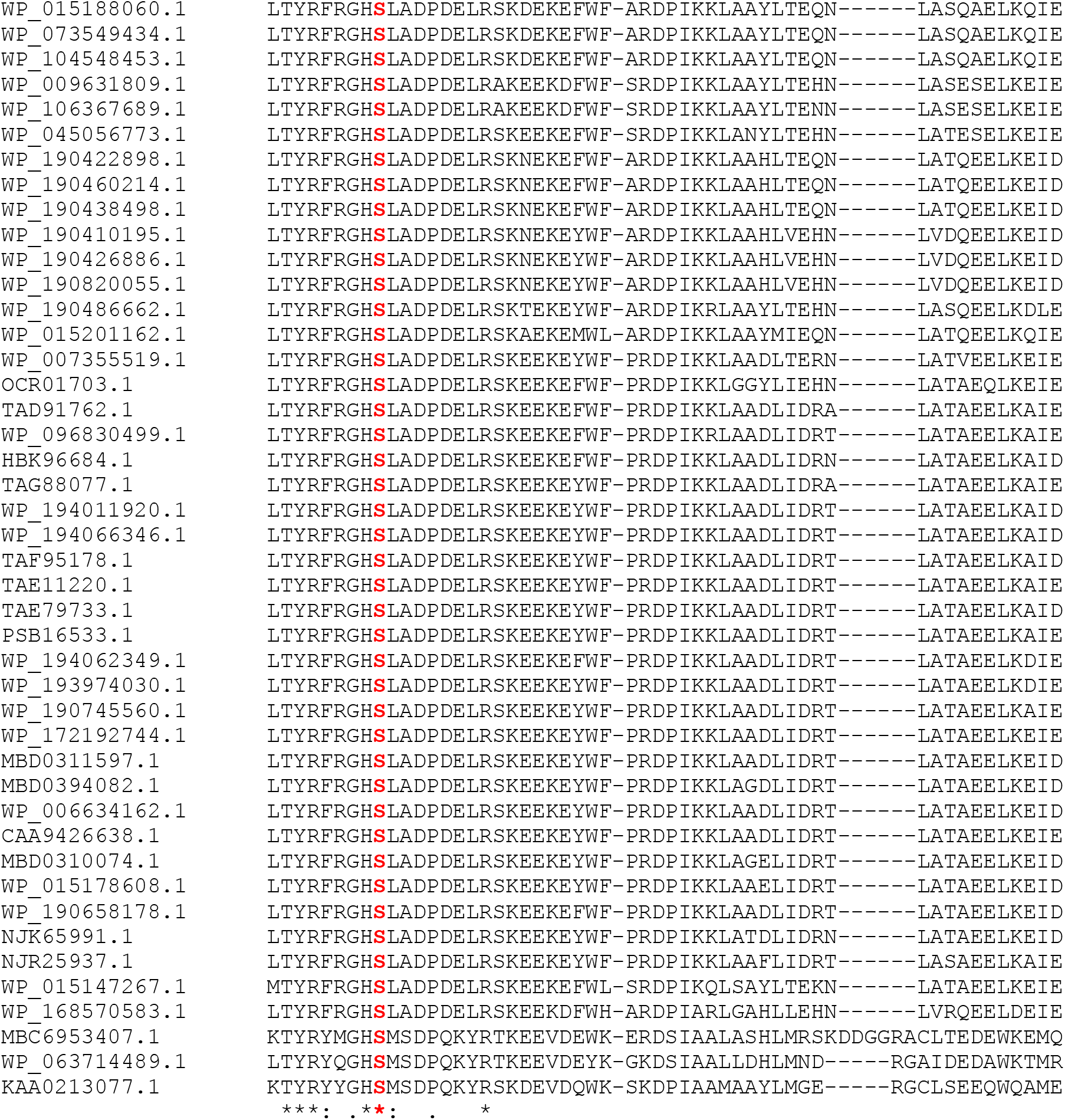
Sequence alignment by ClustalW of 932 cyanobacterial PdhA sequences extracted from Genbank by a blast search on January 11^th^ 2021. The region containing the conserved serine residue (marked in red) is shown.

### PdhA

**Figure S7:**
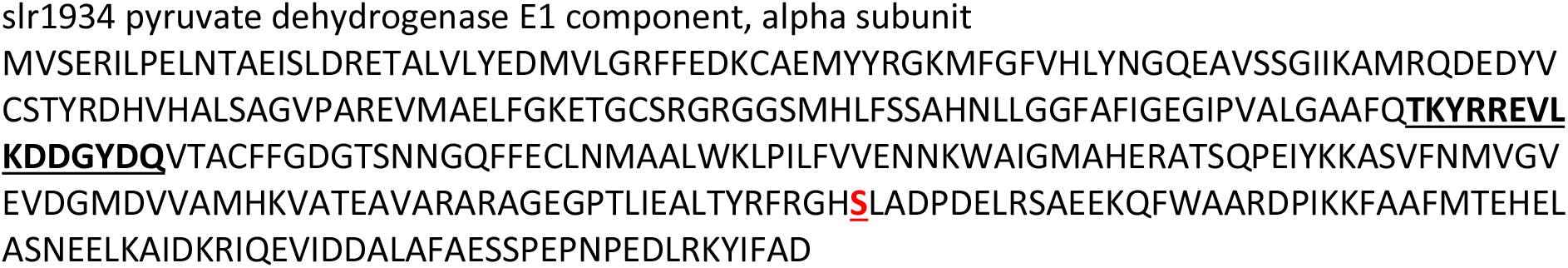
Amino sequence of the PdhA subunit of the PDH complex. The peptide that was used to raise an antibody is underlined. The conserved serine residue is shown in red and underlined as well.

**Figure S8:**
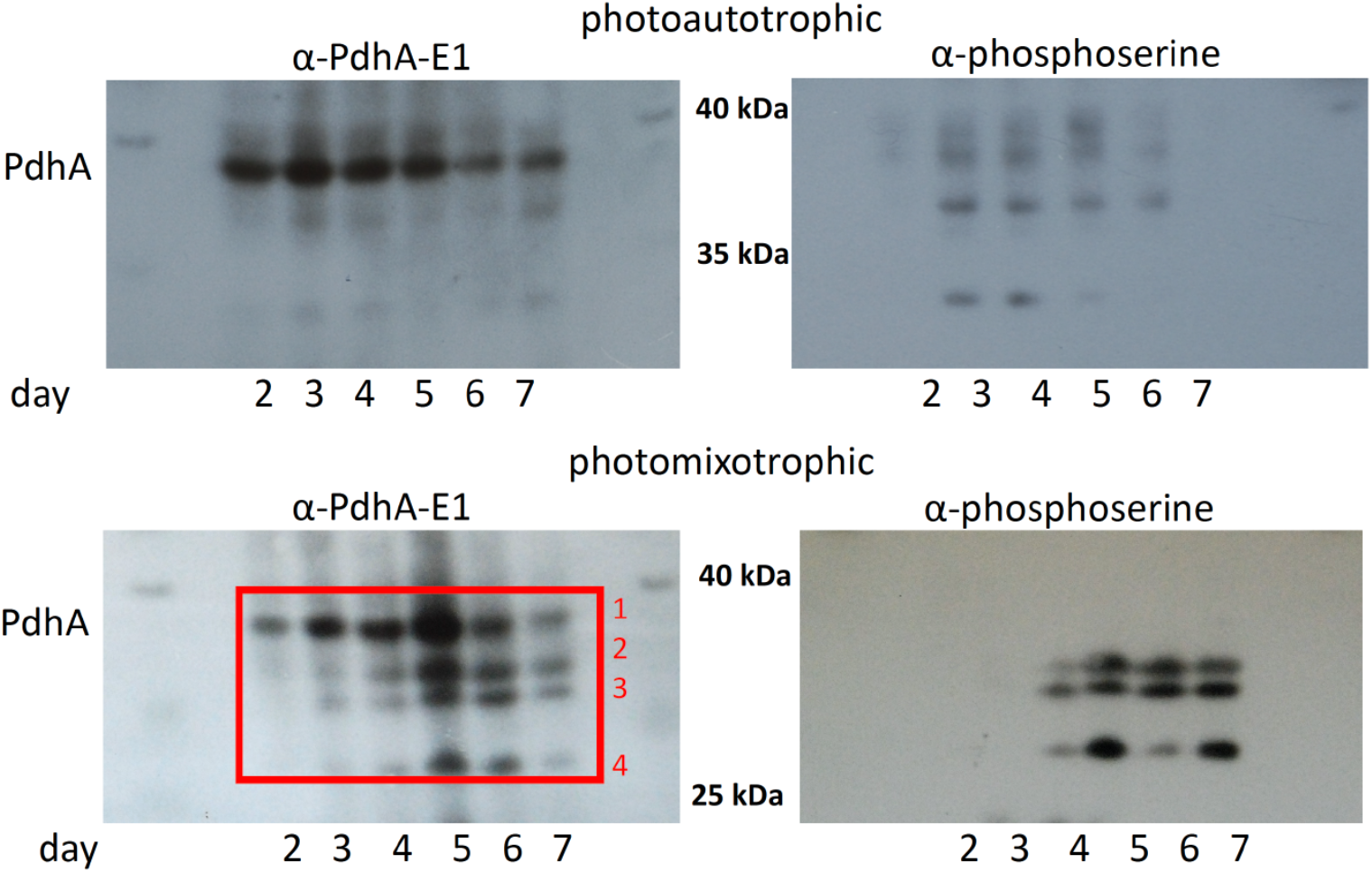
Immunoblots on protein extracts from days 2 to 4 of the wild type grown under photoautotrophic and photomixotrophic conditions. An antibody against the E1 subunit of PdhA of the PDH complex (α-PdhA-E1) as well as an antibody that specifically detects phosphorylated serine residues (α–phosphoserine) were utilized. Representative blots from more than three replicates are shown.

Phosphorylated proteins can be detected by immunoblots due to their modified electrophoretic mobility in contrast to unphosphorylated proteins. Therefore, immunoblots were carried out in order to check if the PDH complex of *Synechocystis* gets phosphorylated under photomixotrophic conditions. Protein extracts were obtained from cells taken from growth experiments on days 2 to 7 as the NADH/NAD^+^ was especially high and as Δ*pfor* displayed its characteristic growth impairment in these days (see Fig. 1A and 1B). An antibody against the E1 subunit of PdhA (α-PdhA-E1) (Fig. S7) and an antibody that binds specifically to phosphorylated serine residues (α-phosphoserine) were used. As we were not able to delete *pdhA* of the PDH complex in *Synechoystis*, we unfortunately could not use a negative control for our immunoblot analyses. However, the expected molecular mass of the E1 subunit of the PDH complex is 38 kDa, and a corresponding strong signal of this size was detected under photoautotrophic and photomixotrophic conditions in the WT by α-PdhA-E1 (Fig. S8).

In line with our hypothesis additional bands appeared on the blots from photomixotrophically grown cells on days 3 to 7 in the range between 26 to 36 kDa. The phosphorylation of a protein results in an additional negative charge which can result in a higher mobility in the SDS gel. All four bands were excised from the gel and analyzed via mass spectrometry. Many peptides which belong to PdhA were detected by MS/MS in the band 1 (red box in Fig. S8) confirming the reliability of the α-PdhA-E1 antibody (Table S3). The MS/MS analyses of the bands 2-4 were aimed to discover the phosphopeptide GHpSLADPDELR in the TiO_2_-enriched peptide fractions. Unfortunately, in contrast to the unmodified peptide, its phosphorylated form remained undetected. These forms were either below detection limits or not present. We subsequently subjected immunoblots as well to a specific antibody against phosphorylated serines (α–phosphoserine). As expected, the phosphoserine antibody did not give a signal for band 1, (which we assumed as being the unphosphorylated form of PdhA) but precisely detected those bands 2-4 that appeared in addition on the blot from photomixotrophically grown cells (which we assumed as being the phosphorylated form of PdhA). Based on these results we cannot unambiguously establish if PdhA of the PDH complex in *Synechocystis* gets phosphorylated. It might as well be that the protein indeed gets phosphorylated and degraded thereupon, which would explain the additional bands that appear under photomixotrophic conditions. In this case the phosphorylation might have been lost in the degraded enzymes. The immunoblots thus give some convincing hints for either a phosphorylation and/or degradation of the PdhA protein in those days in which the NADH/NAD^+^ ratios are high and PFOR gets important for optimal photomixotrophic growth.

**Figure S9:**
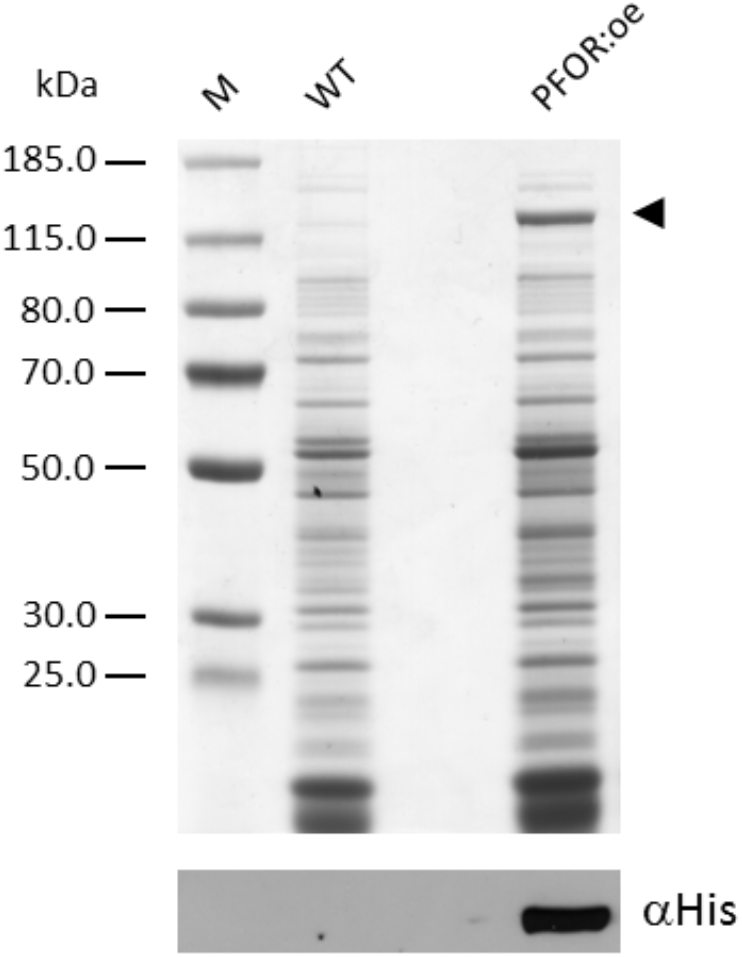
SDS PAGE analysis followed by immunoblotting of *Synechocystis* soluble extracts. Soluble extracts for the wild type (WT) and the mutant overexpressing PFOR (PFOR:oe) containing 15 µg of protein were loaded per lane. The arrowhead indicates the position of over-expressed PFOR.

**Figure S10:**
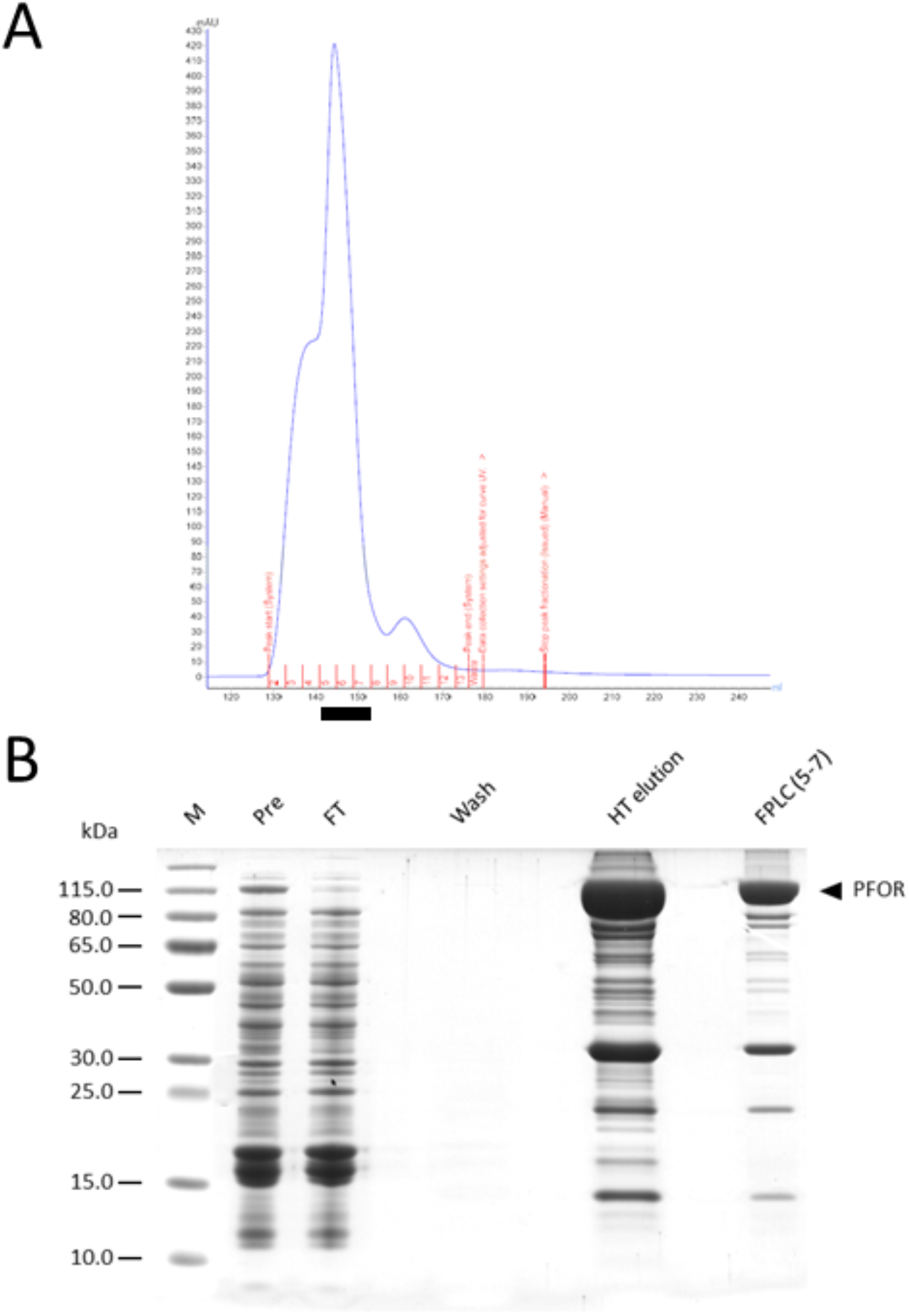
Large-scale PFOR purification. (A) The chromatogram of the FPLC size exclusion run. The collected fractions (5 to 7) are marked by the black bar underneath. (B) Various fractions from the purification procedure were analyzed by SDS PAGE. Soluble extracts before (Pre) and after (Post) the incubation with Talon Cobalt resin, a wash fraction, the His-tag elution and the pooled FPLC fraction (5 t to 7) were loaded on the gel.

**Figure S11:**
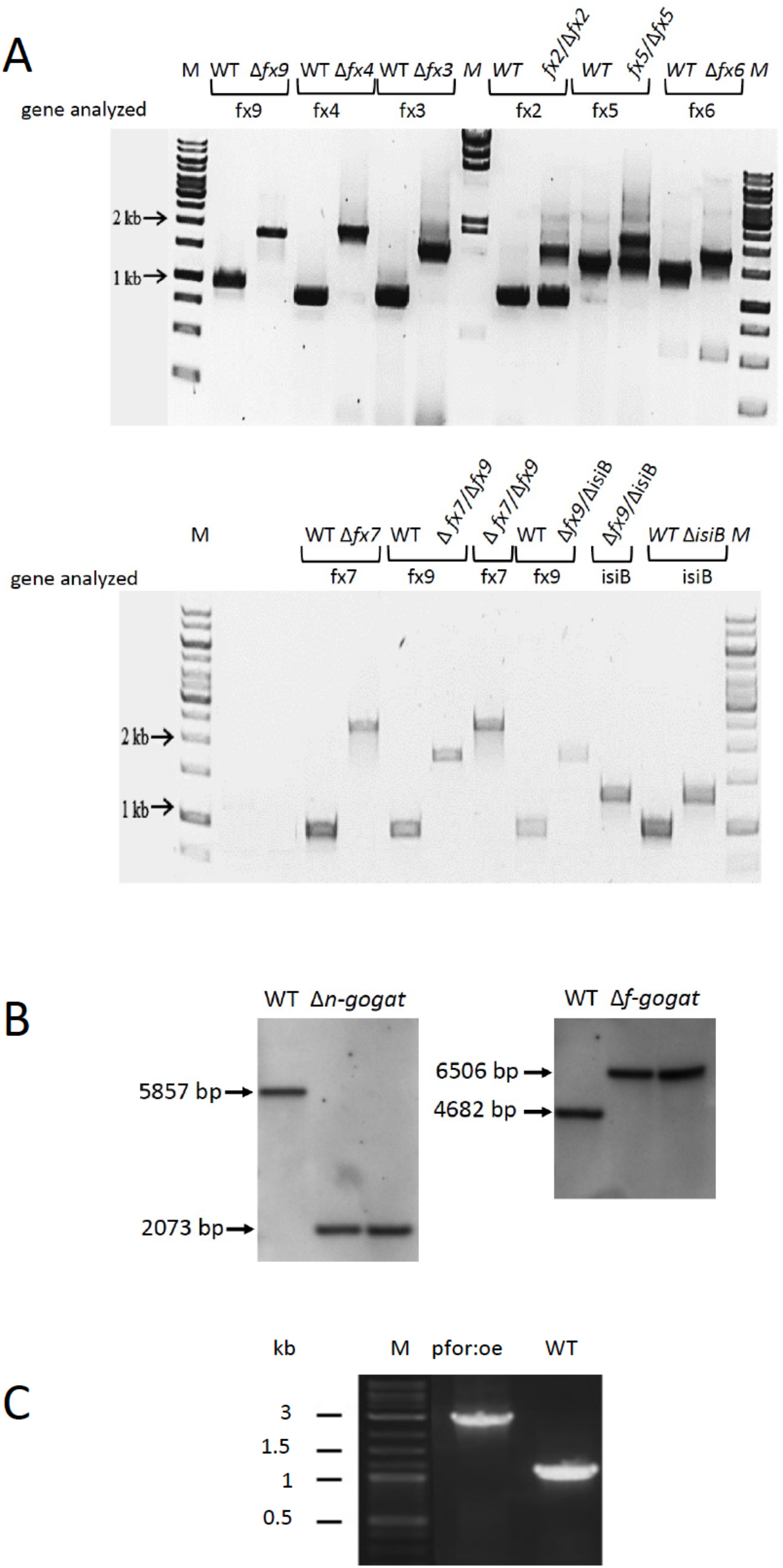
Examination of segregation of mutant strains. (A) PCR analysis of WT, ferredoxin (fx) and flavodoxin (isiB) mutants as indicated. (B) Southern blot of WT and Δ*n-gogat* and Δ*f-gogat* deletion mutants. WT DNA and DNA of two different mutant clones were applied after HindIII digestion. The sizes of the bands are indicated and correspond to those expected due to the mutation. C) PCR analysis of PFOR overexpression (pfor:oe) mutant and WT.

**Figure S12:**
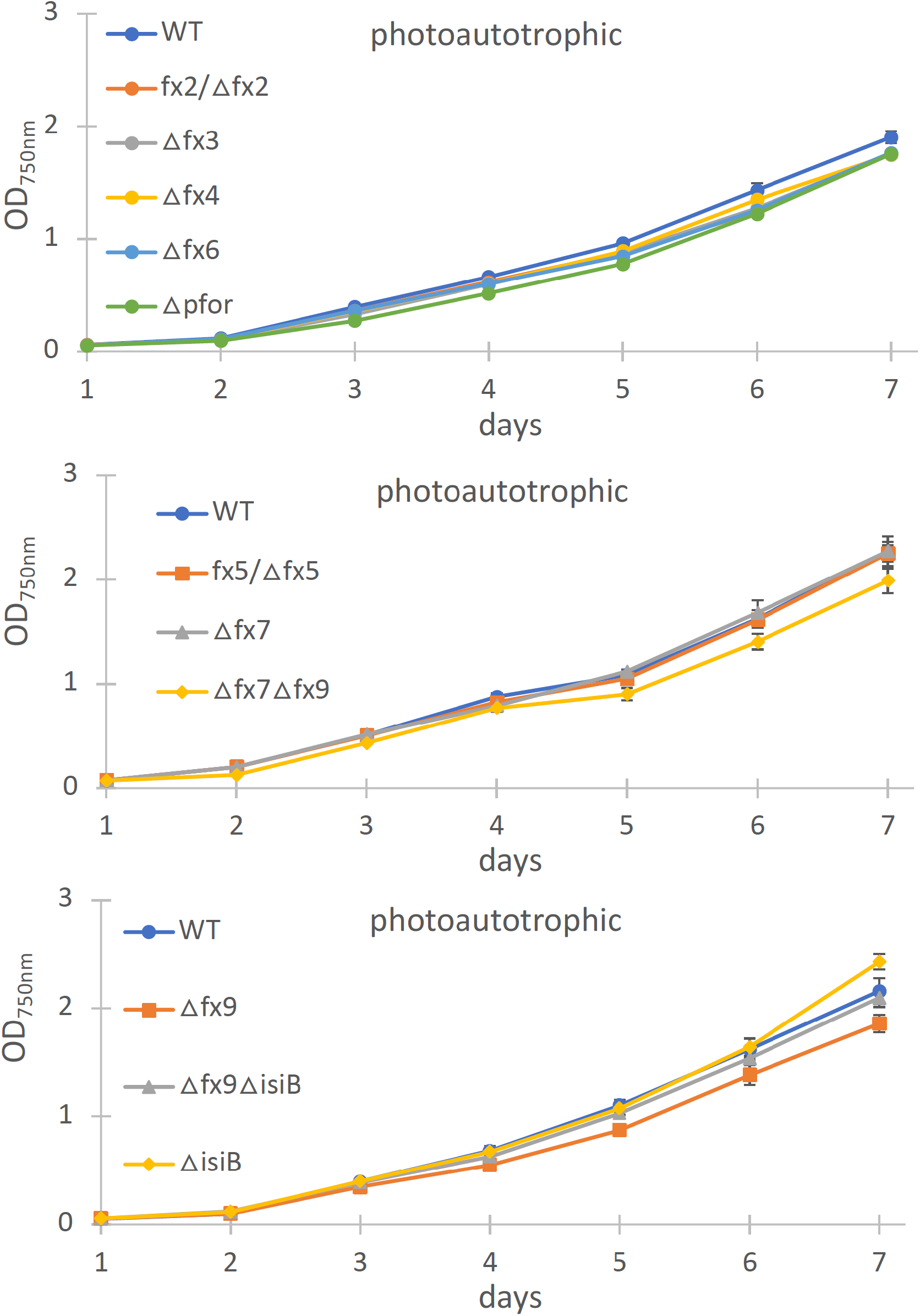
Photoautotrophic growth of different ferredoxin (fx) and the flavodoxin (isiB) deletion mutant as indicated in comparison to the wild type (WT). Shown are mean values ± SD from at least 3 replicates.

**Table S1:**
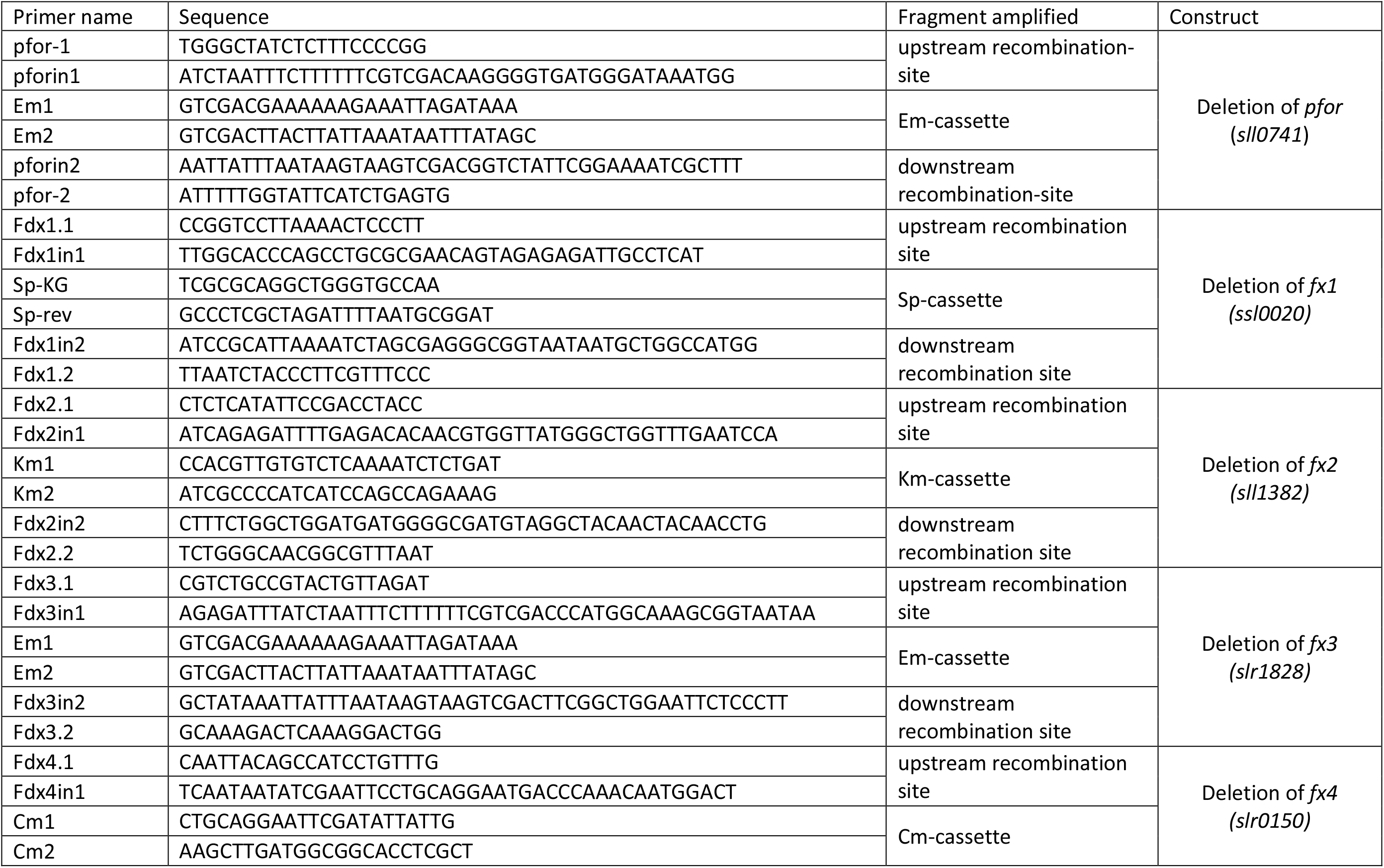
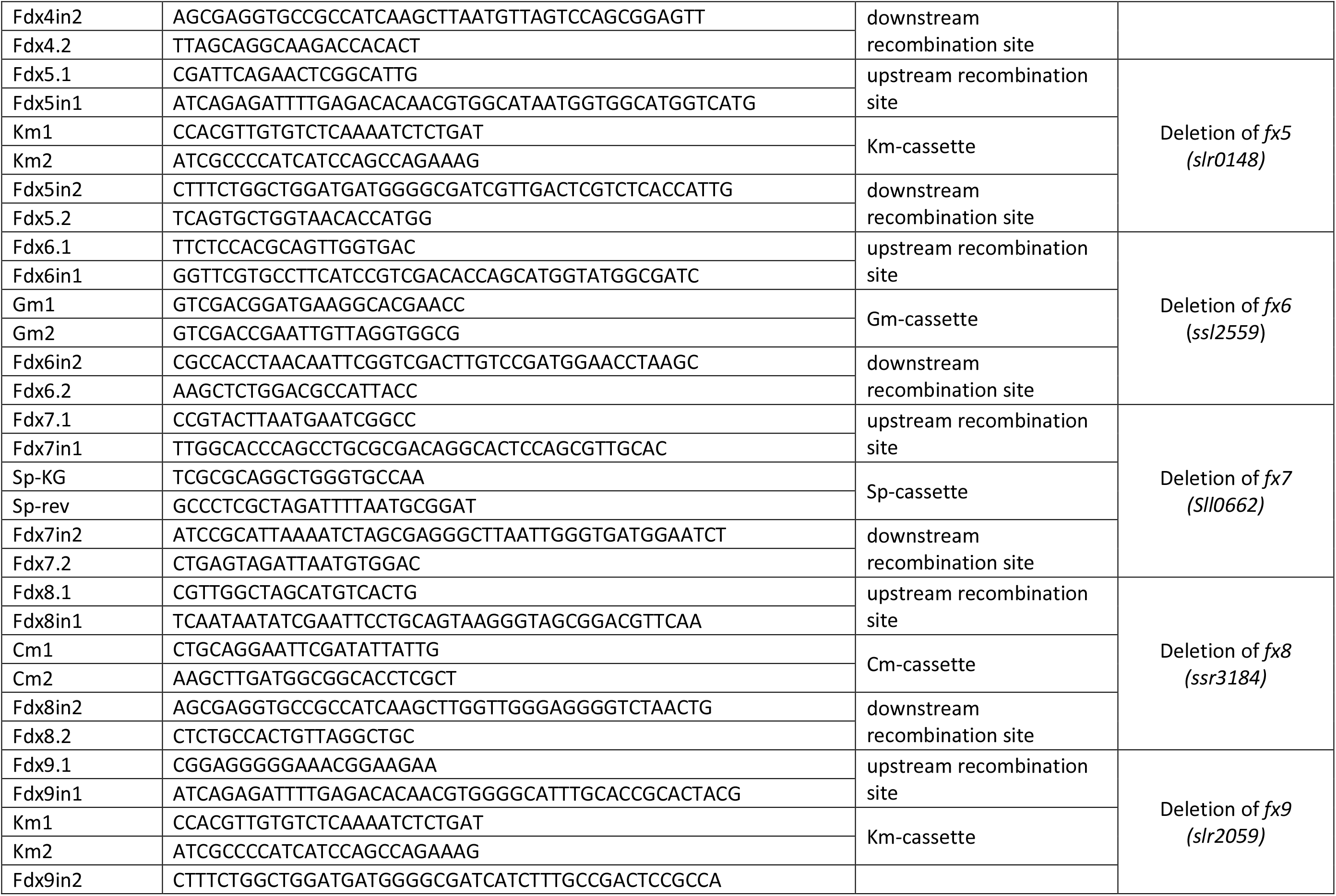
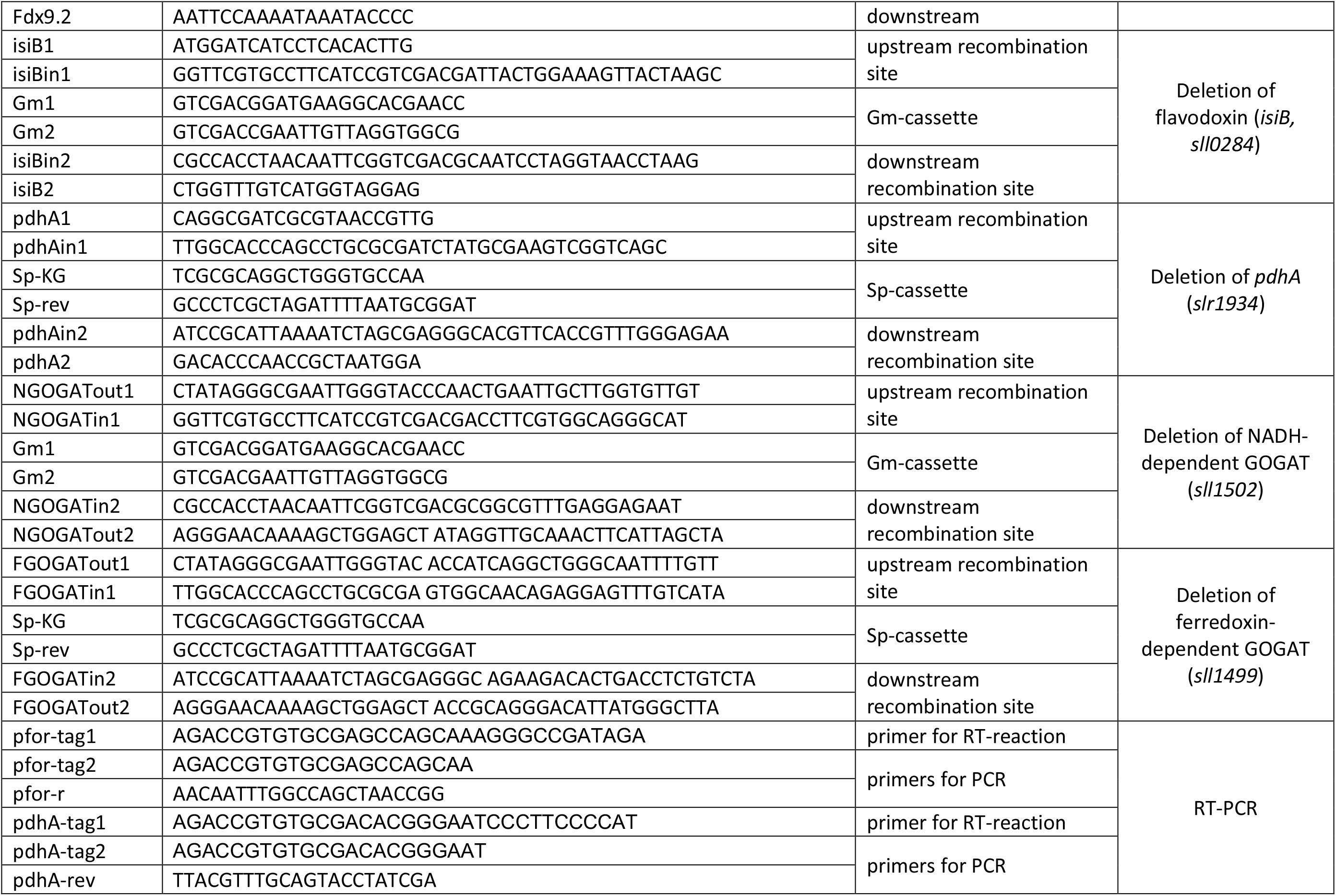
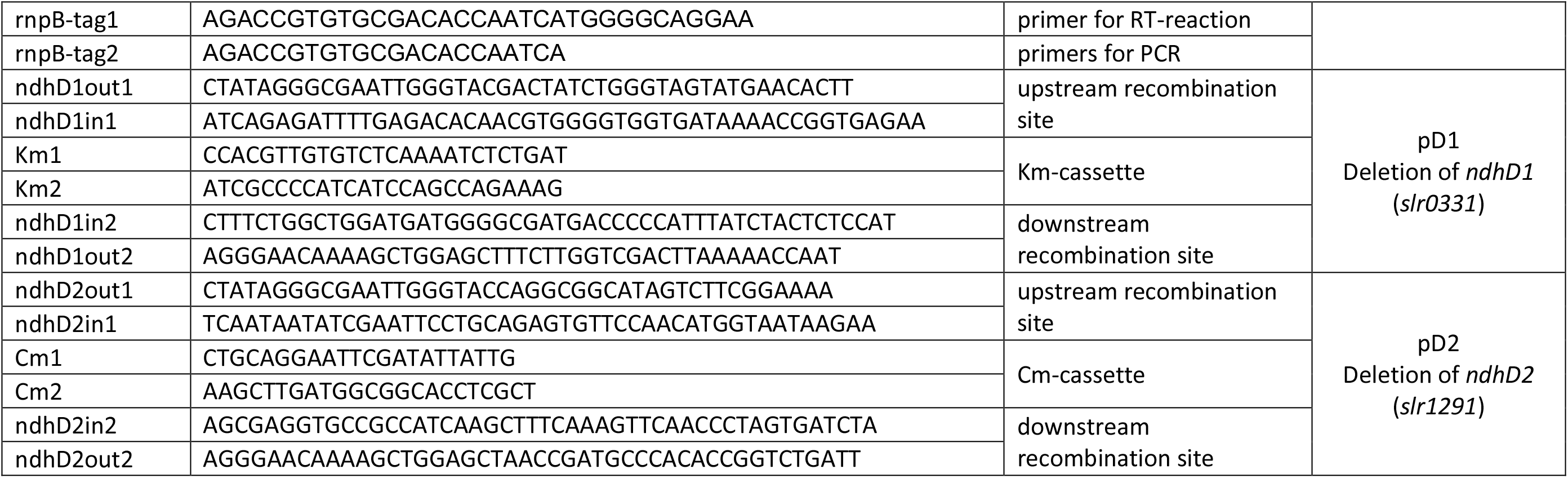
List of primers used in this study to generate deletion strains and for RT-PCR.

**Table S2:** Liste of *Synechocystis* strains and mutants used in this study

| Strain | Marker of genotype | *Synechocystis* WT background | Reference |
| --- | --- | --- | --- |
| WT |  | non-motile GT strain | Trautmann et al. 2012 |
| WTm |  | motile PCC-M strain | Trautmann et al. 2012 |
| ΔspkC | slI0599::km<sup>R</sup> | motile PCC-M strain | This study |
| ΔspkF | slr1225::km<sup>R</sup> | motile PCC-M strain | This study |
| ΔspkI | slI1770::km<sup>R</sup> | motile PCC-M strain | This study |
| ΔspkK | slr1919::km<sup>R</sup> | motile PCC-M strain | This study |
| ΔspkL | slI0095::km<sup>R</sup> | motile PCC-M strain | This study |
| ΔspkB | slr1697::km<sup>R</sup> | non-motile GT strain | France/ Bedu, Marseille, Rippka et al 1979 |
| ΔspkD | slI0776::gm<sup>R</sup> | non-motile GT strain | France/ Bedu, Marseille, Rippka et al 1979 |
| ΔspkE | slr1443::km<sup>R</sup> | non-motile GT strain | France/ Bedu, Marseille, Rippka et al 1979 |
| ΔspkG | slr0152::spec<sup>R</sup>::strep<sup>R</sup> | non-motile GT strain | France/ Bedu, Marseille, Rippka et al 1979 |
| fx2/Δfx2 | slI1382::km<sup>R</sup> | non-motile GT strain | Gutekunst et al. 2014 |
| Δfx3* | slr1828::em<sup>R</sup> | non-motile GT strain | Gutekunst et al. 2014 |
| Δfx4* | slr0150::cm<sup>R</sup> | non-motile GT strain | Gutekunst et al. 2014<br>*please note that the names of fx3 and fx4 are exchanged in Gutekunst et al. 2014 |
| fx5Δfx5 | slr0148::km<sup>R</sup> | non-motile GT strain | This study |
| Δfx6 | ssl2559::gm<sup>R</sup> | non-motile GT strain | This study |
| Δfx7 | slI0662::spec<sup>R</sup> | non-motile GT strain | This study |
| Δfx9 | slr2059::km<sup>R</sup> | non-motile GT strain | This study |
| ΔisiB | slI0284::gm<sup>R</sup> | non-motile GT strain | Gutekunst et al. 2014 |
| Δfx7Δfx9 | slI0662::spec<sup>R</sup>, slr2059::km<sup>R</sup> | non-motile GT strain | This study |
| Δfx7Δfx8Δfx9 | slI0662::spec<sup>R</sup>, ssr3184::cm<sup>R</sup>, slr2059::km<sup>R</sup> | non-motile GT strain | This study |
| Δfx9ΔisiB | slr2059::km<sup>R</sup>, slI0284::gm<sup>R</sup> | non-motile GT strain | This study |
| Δn-gogat | slI1502::gm<sup>R</sup> | non-motile GT strain | This study |
| $\Delta f\text{-gogat}$ | *slI1499::spec<sup>R</sup>* | non-motile GT strain | This study |
| $\Delta ndhD1\Delta ndhD2$ | *slr0331::km<sup>R</sup>, slr1291::cm<sup>R</sup>* | non-motile GT strain | This study |
| $\Delta hk$ | *slI0593::spec<sup>R</sup>* | non-motile GT strain | Theune et al. 2021 |
| $\Delta glgP1\Delta glgP2$ | *slI1356::km<sup>R</sup>, slr1367::spec<sup>R</sup>* | non-motile GT strain | Makowka et al. 2020 |

**Table S3:**
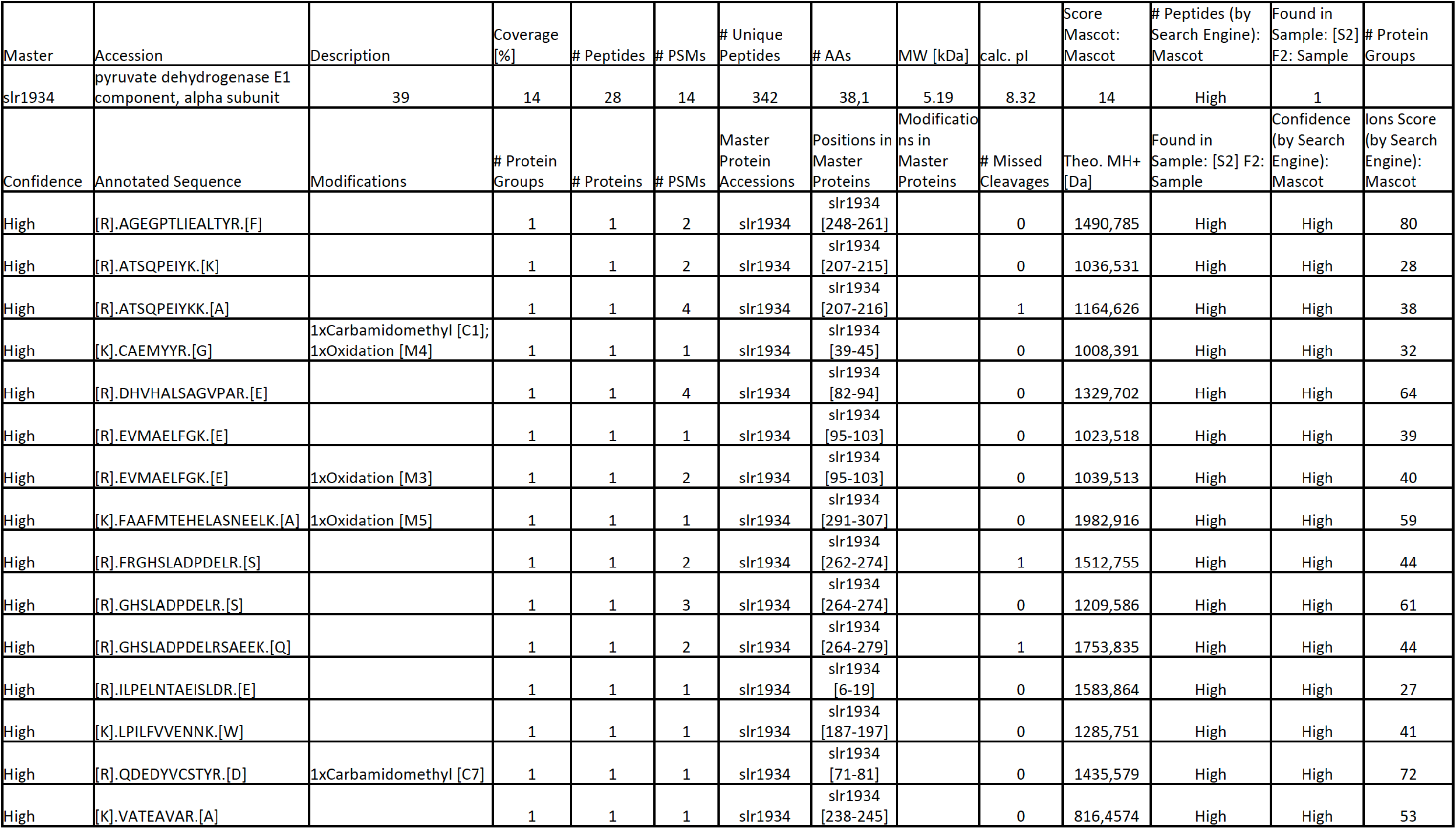
Peptides of the alpha subunit of the pyruvate dehydrogenase (Slr1934) E1 component detected via MS/MS in band No. 1 (red box) of Fig. S8

